# Human repair-related Schwann cells adopt functions of antigen-presenting cells *in vitro*

**DOI:** 10.1101/2022.03.07.483322

**Authors:** Jakob Berner, Tamara Weiss, Helena Sorger, Fikret Rifatbegovic, Max Kauer, Reinhard Windhager, Alexander Dohnal, Peter F. Ambros, Inge M. Ambros, Peter Steinberger, Sabine Taschner-Mandl

**Affiliations:** CCRI, St. Anna Children’s Cancer Research Institute, Vienna, Austria; Department of Plastic, Reconstructive and Aesthetic Surgery, Medical University of Vienna; St. Anna Children’s Hospital, Vienna, Austria; University for Veterinary Medicine, Vienna, Austria; Department of Orthopedics and Trauma Surgery, Medical University of Vienna, Vienna, Austria; Institute of Immunology, Medical University of Vienna, Vienna, Austria

**Author notes:** These authors contributed equally (Jakob Berner, Tamara Weiss). **Corresponding author information:** Sabine Taschner-Mandl, CCRI, St. Anna Children’s Cancer Research Institute, Zimmermannplatz 10, 1090 Vienna, Austria.

**Keywords:** Schwann cell, immunocompetence, PD-L1, immunoregulatory, inflammation, nerve injury, neuropathies, antigen presenting cell

## Abstract

The plastic potential of Schwann cells (SCs) is increasingly recognized to play a role after nerve injury and in diseases of the peripheral nervous system. In addition, reports on the interaction between SCs and immune cells indicate their involvement in inflammatory processes. However, data about the immunocompetence of human SCs are primarily derived from neuropathies and it is currently unknown whether SCs directly regulate an adaptive immune response after nerve injury.

Here, we performed a comprehensive analysis of the immunomodulatory capacities of human repair-related SCs (hrSCs), which recapitulate SC response to nerve injury *in vitro*. We used our previously established protocol for the culture of primary hrSCs from human peripheral nerves and analyzed the transcriptome, secretome, and cell surface proteins for signatures and markers relevant in innate and adaptive immunity, performed phagocytosis assays, and monitored T-cell subset activation in co-cultures with autologous human T-cells.

Our findings show that hrSCs are highly phagocytic, which is in line with high MHCII expression. In addition, hrSCs express co-regulatory molecules, such as CD40, CD80, B7H3, CD58, CD86, HVEM, release a plethora of chemoattractants, matrix remodelling proteins and pro- as well as anti-inflammatory cytokines, and upregulate the T-cell inhibiting PD-L1 molecule upon pro-inflammatory stimulation with IFNγ. Furthermore, hrSC contact reduced the number and activation status of allogenic CD4+ and CD8+ T-cells.

This study demonstrates that hrSCs possess features and functions typical for professional antigen presenting cells *in vitro*, and suggest a new role of these cells as negative regulators of T-cell immunity during nerve regeneration.

**Main points:**

- Human repair-related Schwann cells (hrSC) function as professional antigen presenting cells.
- HrSCs up-regulate PD-L1 upon pro-inflammatory IFNγ stimulation.
- HrSCs hamper CD4+ and CD8+ T-cell activation.

**Graphical abstract:** 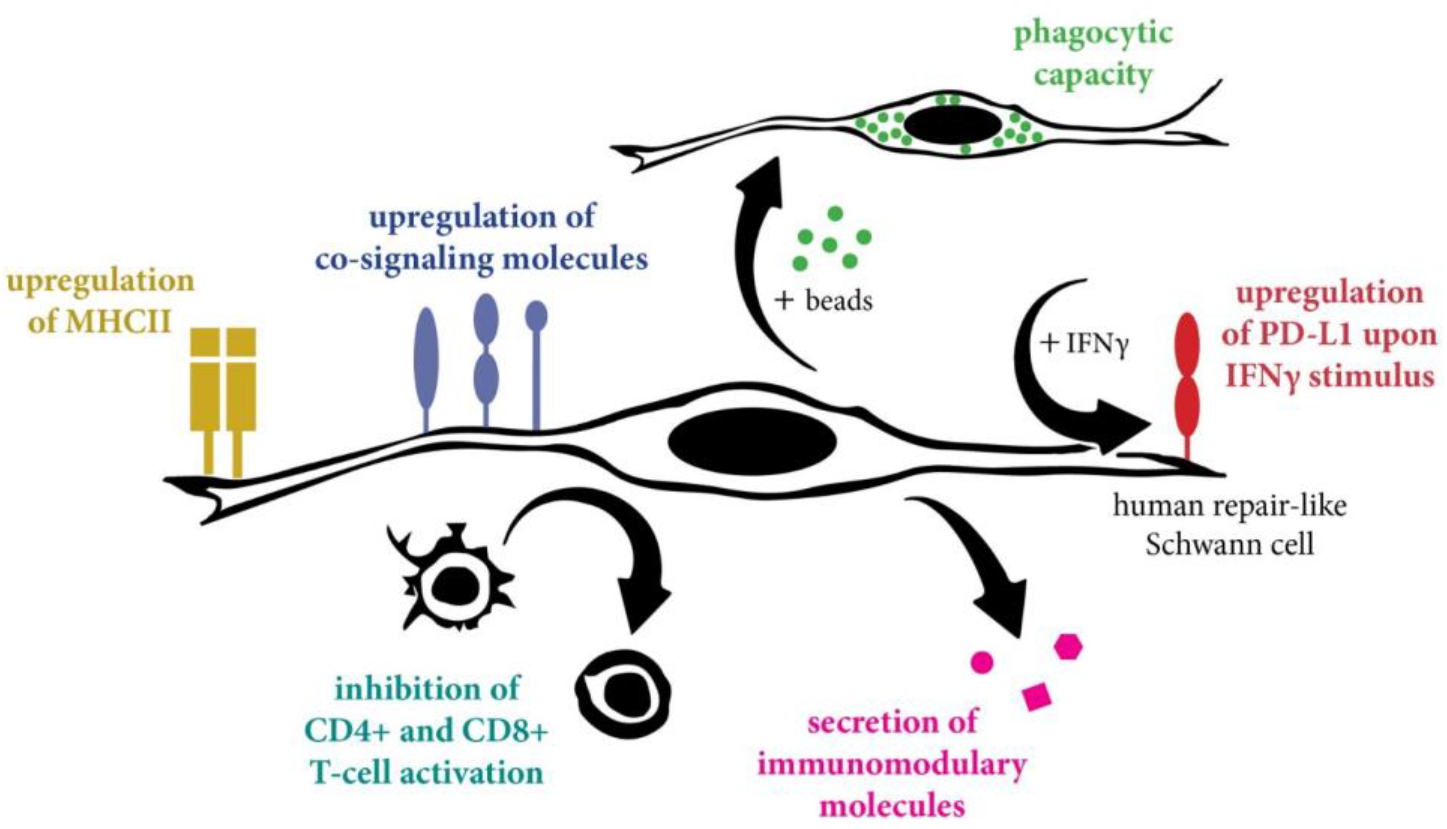

## Introduction

Schwann cells (SCs) are glial cells of the peripheral nervous system and possess capacities that go far beyond the preservation of axon integrity. Upon nerve injury, SCs undergo extensive morphological and expression changes and acquire distinct repair features in a process referred to as ‘adaptive cellular reprogramming’ (Jessen & Mirsky, 2016). In this dedicated repair cell state, SCs re-enter the cell cycle and execute specialized functions to coordinate the multi-step process of nerve regeneration, such as the recruitment of immune cells, the breakdown of myelin debris, remodeling of the extracellular matrix, and the expression of neurotrophic and neuritogenic factors for axon survival, regrowth, and guidance (Gomez-Sanchez et al., 2015; Jang et al., 2016; Jessen & Mirsky, 2016; Nocera & Jacob, 2020; Tofaris, Patterson, Jessen, & Mirsky, 2002; Weiss et al., 2016). Moreover, numerous studies support that the highly adaptive cellular state of SCs plays a role in pathological conditions such as neuropathies and tumor development (Azam & Pecot, 2016; Bunimovich, Keskinov, Shurin, & Shurin, 2017; Direder et al., 2021; Weiss et al., 2021). We have recently shown that tumor-associated SCs in neuroblastic tumors adopt a similar phenotype as upon nerve injury and exert anti-proliferative and pro-differentiating effects through the release of until then unknown neurotrophins, such as EGFL8 (Ambros et al., 1996; Crawford et al., 2001; Direder et al., 2021). As knowledge on the involvement of SCs during regeneration and pathologies is continuously expanding, their immunomodulatory potential gains increasing interest (Armati, Pollard, & Gatenby, 1990; Hörste, Hu, Hartung, Lehmann, & Kieseier, 2008; Zhang et al., 2020). SCs have been demonstrated as immune competent cells that contribute to inflammatory and hereditary neuropathies (Ydens et al., 2013). However, less is known about the impact of human SCs on the inflammatory processes during peripheral nerve regeneration (Bergsteinsdottir, Kingston, Mirsky, & Jessen, 1991; Rutkowski et al., 1999; Toews, Barrett, & Morell, 1998; Weiss et al., 2016).

Similar to any injury site in the body, injured nerves experience an early pro-inflammatory response by the influx of immune cells that is followed by termination of the immune response to allow tissue regeneration. Previous studies showed that SCs secrete a variety of cytokines and chemokines which attract monocytes and neutrophils to the site of nerve injury (Bergsteinsdottir et al., 1991; Rutkowski et al., 1999; Tofaris et al., 2002). Their expression could be partially mediated by axon derived molecules recognized by SCs via toll-like receptors (TLRs) (Goethals, Ydens, Timmerman, & Janssens, 2010; Kaisho & Akira, 2000; Karanth, Yang, Yeh, & Richardson, 2006; Lee et al., 2006; Meyer Zu Horste et al., 2010; Meyer zu Hörste, Hu, Hartung, Lehmann, & Kieseier, 2008). Within injured nerves, recruited and/or tissue resident macrophages adapt a specialized regenerative phenotype with neuroprotective capacities and express proteins associated with an anti-inflammatory profile (Gaudet, Popovich, & Ramer, 2011; La Fleur, Underwood, Rappolee, & Werb, 1996; Ydens et al., 2012). Of note, SCs might be involved in polarizing macrophages towards a regenerative phenotype, but so far, the underlying factors remain unknown (Stratton & Shah, 2016; Stratton et al., 2016).

SCs can also interact with T-cells by expressing major histocompatibility complex class II (MHCII) receptors and co-signaling molecules (Armati et al., 1990; Hörste et al., 2008; Murata & Dalakas, 2000). However, upregulation of MHCII on SCs was primarily reported in neuropathies (Mancardi et al., 1988; Meyer Zu Horste et al., 2010; Van Rhijn, Van den Berg, Bosboom, Otten, & Logtenberg, 2000) and upon treatment with IFNy (Armati et al., 1990; Lilje & Armati, 1997; Samuel, Mirsky, Grange, & Jessen, 1987), which is a potent inducer of MHCII expression in antigen presenting cells (APCs). In contrast, our previous research showed that human repair-related SCs highly upregulate MHCII in culture and within nerve explants independent of IFNy (Weiss et al., 2016). Furthermore, these repair-related SCs expressed genes of co-signaling molecules, MHCII transcriptional co-activator *CIITA*, and other molecules involved in the antigen processing and presentation machinery (Weiss et al., 2016), suggesting a biological relevance of MHCII expressing SCs in response to injury.

APCs function as local modulators of T-cell response upon inflammatory stimulation. The outcome of this modulation is dependent on the expression of sets of co-stimulatory or co-inhibitory surface molecules recognized by T-cells together with MHCII. Indeed, rodent SCs could be induced to activate T-cells by presenting endogenous as well as exogenous antigens (Duan et al., 2007; Kingston et al., 1989; Spierings, De Boer, Zulianello, & Ottenhoff, 2000; Steinhoff & Kaufmann, 1988; Wekerle, Schwab, Linington, & Meyermann, 1986). Moreover, T-cell activation through MHCII expressing SCs has been associated with post-traumatic inflammation and neuropathic pain in diseased peripheral nerves of rodents (Hartlehnert et al., 2017). Hence, the SC function as non-professional APC has mainly focused on the promotion of T-cell activation resulting in (auto-) inflammatory or infectious neuropathies, rather than suppression of activated T-cells. The latter is executed by APCs to restore immune homeostasis and prevent auto-immunity to self-proteins. In line with a potential T-cell inhibiting function of SCs, our previous transcriptomic analyses have indicated that primary human SCs express genes associated with T-cell suppression such as *PD-L1* and *DC-HIL* (Weiss et al., 2016).

Based on the increasing body of studies supporting the immunocompetence of SCs and their recognized role in nerve injury and disease (Meyer zu Hörste et al., 2008; Weiss et al., 2021; Zhang et al., 2020), we here set out to investigate immunoregulatory features of human SCs in an injury condition. To this end, we cultured primary human SCs and performed phagocytosis assays, analyzed the secretion of immunomodulatory mediators, and profiled their repertoire of co-signaling molecules as well as upon TLR and inflammatory stimulation. We further assessed the ability of SCs to modulate T-cell activation and polarization *in vitro*.

## Methods

### Human material

The collection and research use of human peripheral nerve tissues and human tumor specimen was conducted according to the guidelines of the Council for International Organizations of Medical Sciences and World Health Organisation and has been approved by the local ethics committees of the Medical University of Vienna (EK2281/2016 and 1216/2018). Informed consent has been obtained from all patients participating in this study.

Neuroblastoma cell lines and primary cultures are available upon request. Primary Schwann cell cultures and tumor tissues are limited materials and therefore cannot be provided.

### Isolation of primary human Schwann cells

SCs were isolated, cultured and enriched as previously described (Weiss, Taschner-Mandl, Ambros, & Ambros, 2018). Briefly, peripheral nerves were cut into 2-3 cm pieces and nerve fascicles were pulled out of the surrounding epineural tissue. The isolated fascicles were cut into ~0.5 cm pieces and incubated in a digestion solution containing αMEM GlutaMAX^TM^ (Gibco), 10% FCS (PAA), 1% Pen/strep (Pan Biotech), 1 mM sodium pyruvate (Pan Biotech), 25 mM HEPES (Pan Biotech), 0.125% collagenase Type IV (Gibco), 1.25 U/mL Dispase II (Sigma-Aldrich) and 3 mM CaCl (Sigma-Aldrich) at 37 °C, for 20 h. The digested tissue was pelleted and resuspended in SC expansion medium (SCEM) containing MEMα, 1% Pen/Strep, 1 mM sodium pyruvate, 25 mM HEPES, 10 ng/mL hu FGF basic (PeproTech), 10 ng/mL hu Heregulin-β1 (PeproTech), 5 ng/mL hu PDGF‐AA (PeproTech), 0.5% N2 supplement (Gibco), 2 μM forskolin (Sigma-Aldrich) and 2% FCS. Cells were seeded in 0.01% Poly-L-lysine (PLL, Sigma-Aldrich) and 4 μg/ml laminin (Sigma-Aldrich) coated culture dishes. Half of the medium was changed twice a week. As passage 0 (p0) cultures consisted of SCs and fibroblast-like cells, SCs were enriched before experimentation, by exploiting their differential adhesion potential to plastic, described in (Weiss et al., 2018). As previously shown, human SCs adopt a repair-related phenotype in culture (Weiss et al., 2016) and SCs are referred to as human repair-related SCs (hrSC).

### Neuroblastoma cell lines

The used neuroblastoma cell lines (NB cells) are derived from biopsies or surgical resection of aggressively behaving, high-risk neuroblastomas. In-house established, low passage NB cell lines STA-NB-6, 7, −10 and −15 as well as the cell lines, SH-SY5Y, IMR5 and CLB-Ma (kindly provided by Dr Valerie Combaret, Centre Leon Berard, France) (I M Ambros et al., 1997; Biedler, Helson, & Spengler, 1973; Biedler, Roffler-Tarlov, Schachner, & Freedman, 1978; Combaret et al., 1995; Fischer & Berthold, 2003; Momoi, Kennett, & Glick, 1980; Stock et al., 2008) were used for experimentation and cultured in MEMα GlutaMAX^TM^, 1% Pen/Strep, 1 mM sodium pyruvate, 25 mM HEPES and 10% FCS. NB cells are used in this study as model to reflect neuronal cells.

### Primary human T-cells

For the T-cell isolation, a buffy coat was obtained from the Austrian Red Cross and diluted 1:4 in 1x PBS. Density gradient centrifugation was performed by transferring the blood onto 20 mL Lymphoprep solution (StemCell Technologies) and centrifugation for 30 min at 400g at room temperature (RT) without breaks. Mononuclear cells were carefully removed from the interphase layer and transferred into 50 mL 1x PBS and centrifuged at 300g for 10 min. Then, the medium was removed and T-cells were isolated with the Pan T-cell Isolation Kit (Miltenyi Biotec) according to the manufacturer’s protocol using magnetic activated cell sorting (MACS). Briefly, cells were counted and resuspended in 40 μL of MACS buffer (PBS, pH 7.2, 0.5% bovine serum albumin (BSA), 2 mM EDTA) per 10^7^ cells. For each 40 μL, 10 μL of PAN T-Cell Biotin Antibody Cocktail (Miltenyi Biotec) was added and incubated for 5 minutes at 4°C. Then 30 μL of MACS buffer and 20 μL of Pan T-Cell Microbead Cocktail was added per each 50 μL solution and incubated for additional 10 min at 4°C. For the magentic separation, MACS LS columns (Miltenyi Biotec) were placed in the magnetic field of a MACS separator (Miltenyi Biotech) and the column was rinsed with 3 mL MACS buffer. The cell suspension was added and the flow-through containing the unlabelled, CD3 positive T-cells, was collected. Cells were frozen in Cryostor freezing medium (Biolife Solutions) and stored in liquid nitrogen until the day of the experiments.

### RNA Sequencing and data analysis

RNA-sequencing datasets have been previously published and are available at the Gene expression omnibus (GEO) repository under the identifiers GSE94035 (MNC, n=5), GSE90711 (SC, n=5), GSE90711 (NB primary cultures, n=5 over 3 patient cultures STA-NB-6, STA-NB-7 and STA-NB-15). RNA isolation, library preparation and sequencing on a Illumna Hiseq 2000 platform were performed as previously described (Weiss et al., 2016, 2021). Short read sequencing data was quality checked using FASTQC (http://www.bioinformatics.babraham.ac.uk/projects/fastqc) and QoRTs (Hartley & Mullikin, 2015) and then aligned to the human genome hs37d5 (ftp://ftp.1000genomes.ebi.ac.uk/) using the STAR aligner (Dobin et al., 2013) yielding a minimum of 11.6 million aligned reads in each sample. Further analysis was performed in R statistical environment using Bioconductor packages (Gentleman et al., 2004). Count statistics for Ensembl (GRCh37.75) genes were obtained by the “featureCounts” function (package “Rsubread”) and differential expression analysis was performed by edgeR and voom (Law, Chen, Shi, & Smyth, 2014; Ritchie et al., 2015). For differential gene expression analysis only genes passing a cpm (counts per gene per million reads in library) cut-off of 1 in more than two samples were included. All p-values were corrected for multiple testing by the Benjamini-Hochberg method. Genes with an adjusted q-value <0.05 and a log2 fold change > 1 (|log2FC|>1) were referred to as ‘significantly regulated’ and used for functional annotation analysis via gene set enrichment analysis (GSEA) using MSigDB according to Subramanian, Tamayo, et al. (Subramanian et al., 2005) and Mootha, Lindgren, et al. (Mootha et al., 2003)

### Phagocytosis assay

5×10^4^ enriched p1 hrSCs were seeded per well of an 8‐well chamber slide (Ibidi) coated with PLL/laminin and cultured in SCEM. After 48 h, half of the medium was replaced with fresh SCEM containing 1 μm big carboxylate‐modified polystyrene, fluorescent yellow‐green latex beads (SIGMA-Aldrich) at a concentration of 8×10^6^ beads/well (∼100 beads/cell) for 15 h at 37°C. Thereafter, cells were washed three times with 1x PBS and fixed with Roti‐Histofix (Roth) for 10 min at RT. Cells were stored at 4°C in 1x PBS until multi-color immunofluorescence staining was performed.

### Immunofluorescence stainings

All antibody details, dilutions and incubation times are listed in **Supplementary Table 1**. If not stated otherwise, the staining procedure was performed on RT and a washing step (3 times with 1x PBS for 5 min) was performed after each antibody incubation step, except after permeabilization. For extracellular staining, grown cells were blocked with 1x PBS containing 3% goat serum (DAKO) for 30 min at RT, followed by incubation with antibodies against extracellular targets diluted in 1x PBS containing 1% BSA (Sigma-Aldrich) and 1% goat serum. Cells were then incubated with secondary antibodies diluted in 1x PBS containing 1% BSA and 1% goat serum. For permeabilization, cells were exposed to 1x PBS containing 1% BSA, 0.3% Triton-X (Sigma-Aldrich) and 3% goat serum for 10 min. Thereafter, cells were incubated with primary antibodies against intracellular targets diluted in 1x PBS containing 1% BSA, 0.1% Triton-X and 1% goat serum, followed by incubation with secondary antibodies diluted in 1X PBS containing 1% BSA, 0.1% Triton-X and 1% goat serum. Afterwards, 2 μg/mL 4′,6-Diamidin-2-phenylindol (DAPI, Sigma-Aldrich) in 1X PBS was added for 2 min followed by a final washing step. Cells were embedded in Fluoromount-G^TM^ mounting medium (Southern Biotech) and stored at 4°C. Images were taken with a confocal laser scanning microscope (Leica Microsystems, TCS SP8X) using Leica application suite X version 1.8.1.13759 or LAS AF Lite software. Confocal images are depicted as maximum projection of total z-stacks and brightness and contrast were adjusted in a homogenous manner using the Leica LAS AF software.

### FACS characterization of hrSCs

All antibodies used for flow cytometry stainings are listed in **Supplementary Table 1**. If not stated otherwise, the staining procedure was performed on 4°C. For all phenotyping experiments, cells were cultured in duplicates in each condition. HrSCs were cultured in the presence of IFNγ (10^3^ U/mL, Bio-Techne Ltd.), LPS (10 ng/mL, Sigma-Aldrich,), Poly:IC (2 μg/mL, Bio-Techne Ltd.) cross-linked CD40L (500 ng/mL, Bio-Techne Ltd.) and IL-1β (10^4^ U/mL Bio-Techne Ltd.) for 24 h. Cells were harvested with Accutase (Sigma-Aldrich) and transferred into FACS tubes containing 200 μL FACS buffer (0.1% BSA and 0.05% natrium acides in 1x PBS). Cells were washed once with FACS buffer at 1200rpm for 5 min, resuspended in 50 μL FACS buffer and incubated with 50 μL of an antibody master mix in FACS buffer for 30 min in the dark. Then, cells were washed with FACS buffer and resuspended in 100 μL of Cytofix/Cytoperm solution (BD Biosciences), incubated for 20 min, and again washed with BD 1x perm buffer (BD Biosciences). Next, cells were resuspended in 100 μL 1x perm buffer containing the S100 antibody and incubated for 30 min in the dark. Cells were then washed in 1x perm buffer and resuspended in 100 μL 1x perm buffer with the secondary antibody for 20 min. After a washing step in 1x perm buffer, cells were washed with FACS buffer and resuspended in 100 μL FACS buffer. All samples were measured with a FACSFortessa flow cytometer equipped with 5 lasers (355, 405, 488, 561 and 640 nm) and the FACSDiva software version 8.0 (BD Biosciences) was used.

### T-cell proliferation assay

For all T-cell experiments, hrSCs and freshly thawed human CD3^+^ T-cells were used in various conditions. For IFNy stimulation, hrSCs were cultured in the presence of 10^3^ U/mL IFNγ for 24 h prior to the T-cell co-culture experiments. For co-culture, p1 hrSCs were harvested, counted and seeded at 4×10^4^ cells per 96 well plates in duplicates. T-cells were thawed, washed once with 1XPBS and centrifuged at 300g for 7 min at RT. Cells were counted and labelled with CFSE (Thermo Fisher) at 1 μL/10^7^ cells for 10 min at 37°C. Thereafter, 1 mL FCS buffer was added for 2 min at RT and then washed with αMEM at 300g for 7 min. Then, cells were FACS sorted for intact cells using a FSC vs SSC gate using the FACS Aria instrument (BD Bioscience). The obtained cells were washed with αMEM at 300g for 7 min, counted and 1×10^5^ cells were seeded to the hrSCs (co-culture) or cultured alone (controls) in 96 well plates. For the T-cell stimulation, 0.25 μL of anti-CD3/CD28 beads (Gibco) were added. Culture medium containing CD3/CD28 beads was thoroughly replenished every 3 days by one half.

Cells were analysed via flow cytometry at different time points, i.e. at day 2, 4 and 10. Therefore, cells were harvested with Accutase and washed once with FACS buffer. All antibodies used for flow cytometry stainings are listed in **Supplementary Table 1**. Extracellular staining was performed with 50 μL of antibody mix containing all extracellular antibodies (CD3-FITC, CD4-PerCP, CD8APC-Cy7, CD25-PE-Cy7) in 50 μL of FACS buffer for 30 min at 4°C. Cells were washed and incubated in Fix/Perm solution (Thermo Fisher) at 4°C for 30 min. For the permeabilization, 1x perm buffer (Thermo Fisher) was added and cells were centrifuged at 300g for 7 min. Then, the supernatant was discarded and the S100 antibody for intracellular staining was added in 100 μL 1x perm buffer and incubated for 20 min at RT in the dark. After this, cells were washed once with 1x perm buffer, once with FACS buffer and resuspended in 100 μL FACS buffer. For exact quantification of absolute cell numbers, 10 μL AccuCheck Counting Beads (LifeTechnologies) were added to each sample prior to FACS analysis at a FACSFortessa flow cytometer. For data analysis, FACSDiva software version 8.0 was used. Gating for CD4^+^ Th subsets was performed in accordance to Mahnke *et al*. 2013 (Mahnke, Beddall, & Roederer, 2013).

### Protein array

The RayBio G-Series Human Cytokine Antibody Array 4000 Kit (RayBiotech Inc.) was used to assay secretomes of cell supernatants pooled from 2 independent experiments each from hrSC (n=5), hrSCs co-cultured with 5 different NB cell cultures (n=5) or NB cell cultures alone (n=5). A total of 274 factors were evaluated (for a complete list of factors, refer to https://www.raybiotech.com/human-cytokine-array-g4000-4/). Arrays were processed according to the manufacturer's instructions. Briefly, protein array membranes were blocked with Blocking Buffer (RayBiotech Inc.) for 30 min at RT. Membranes were then incubated with 100 μL of undiluted sample for 2 h. After extensive washing with Wash Buffer I and II (RayBiotech Inc.) to remove unbound materials, the membranes were incubated with biotin-conjugated antibodies for 2 h at RT. The membranes were then washed and incubated with streptavidin-fluorescin, again for 2 hours at RT, followed by final washing steps. Finally, fluorescence signals were obtained with the GenePix 4000 array scanner (Molecular Devices) using the green channel (Cy3) at an excitation frequency of 532 nm and 700 PMT. The image files generated in this way were aligned to respective .gal files (RayBiotech) and Gene Pix Pro 7 (Molecular Devices) was used to create .gpr files. Each spot was manually inspected on the .gpr file images to ensure accuracy. After background correction and normalization to the internal control, the mean fluorescence intensity (MFI) values were combined for all cell lines and proteins that were differentially expressed (q < 0.05) between hrSCs and hrSCs in co-cultures, compared to neuronal cells as controls, were selected for visualization using the Qlucore Omics Explorer V3.1 software.

### Quantification and statistical analysis

If not mentioned otherwise, statistical analysis was performed with R version 3.4.2 within the R studio interface including publicly available packages CRAN, GGPLOT2, GGBEESWARM and RESHAPE. For pair-wise comparison paired t-tests were used, for multiple comparisons two-way ANOVA using a post-hoc Holm p value correction was used. P values of less than 0.05 were considered significant and displayed as *, p values of less than 0.01 were displayed as **, p values of less than 0.001 were displayed as ***.

### Data and code availability

No original code has been generated in this study. Original/source data for all figures and supplementary figures are available upon request. RNA-sequencing data are available at the Gene Expression Omnibus (GEO) repository (Home - GEO - NCBI (nih.gov) under the identifier GSE94A035, GSE90711 and GSE90711.

## Results

### Human repair-related Schwann cells show a phagocytic capacity

APCs are characterized by their phagocytic ability of exogenous material to process and present antigens via MHCII. To evaluate whether human SCs can take-up material different from myelin, we applied our previously established protocol for the culture of primary SCs from human peripheral nerves (Weiss et al., 2018). As human SCs possess a repair-like phenotype and perform repair-associated functions in culture (Weiss et al., 2016), they are referred to as human repair-related SCs (hrSCs) in the following. The cultured hrSCs showed the typical spindle-shaped morphology with a swirled parallel alignment **(Fig. 1A)** and were characterized by immunostainings for the SC marker NGFR (also known as TNR16 or p75^NTR^) **(Fig. 1B)**. The co-staining for vimentin, an intermediate filament expressed by SCs and fibroblasts, visualized a straight filament network within the long hrSC processes and a more branched appearance in fibroblasts **(Fig. 1B)**. To obtain information about their phagocytic capacity, we challenged the hrSCs with green fluorescent latex beads (1 μm diameter) for 15 hours and stained the cultures for NGFR and vimentin. 3D confocal image analysis demonstrated that hrSCs were able to phagocytose the beads and to accumulate them within the cell body **(Fig. 1C)**.

**Figure 1.**
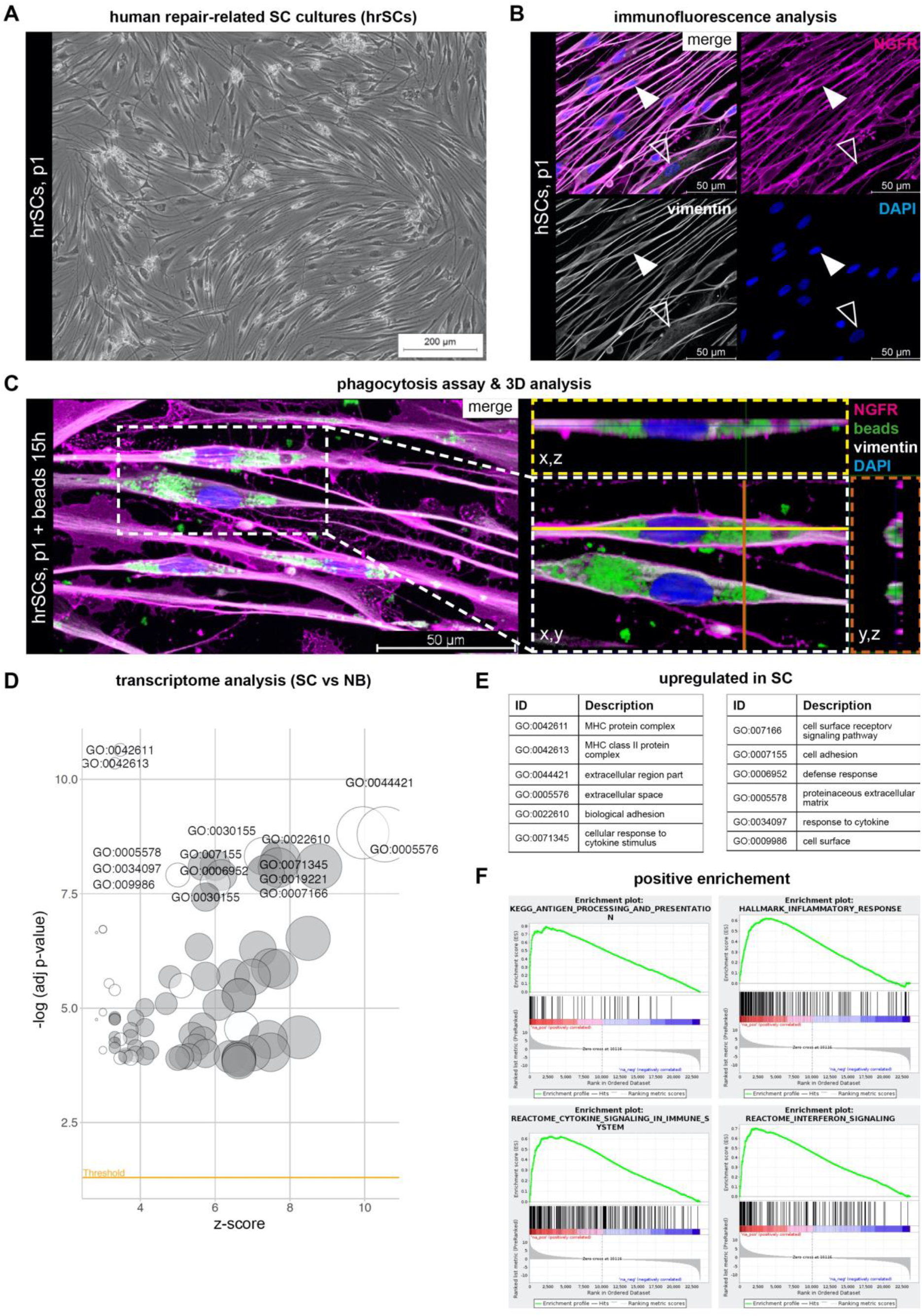
Phagocytosis potential and inflammatory response of hrSCs. **(A)** Phase contrast image of a representative passage 1 (p1) hrSC culture. **(B)** Immunostaining of p1 hrSCs for SC marker NGFR (magenta), intermediate filament vimentin (grey) and nuclear stain DAPI (blue); arrowheads indicate a NGFR negative and vimentin positive fibroblast. **(C)** 3D confocal image analysis of hrSCs exposed to 1 μm in diameter green fluorescent latex beads for 15 hours. Cross sections show internalized beads within the SC cytoplasm. **(D)** Gene ontology (GO) analysis of differentially expressed genes by RNA-seq between hrSCs (n=5) and NB cultures (n=5 independent biological replicates of 3 donors), –log10 of the enrichment p values (cut-off <0.05) for filtered GO categories are plotted relative to Z-scores of average ratios in each category. Circle size represents the fraction of regulated genes per GO term. **(E)** Top 12 GO terms among genes upregulated in hrSCs vs NB cultures. **(F)** Gene set enrichment analysis (GSEA) plots of hrSCs compared to NB cultures. Source data are provided in **Suppl. Tables 2-4**.

### Transcriptome profiling of human repair-related Schwann cells revealed immunomodulatory gene signatures

The phagocytic capacity of hrSCs towards cell-extrinsic material prompted us to evaluate whether pathways associated with inflammation and antigen presentation were active in hrSCs. To this end, we interrogated the transcriptome data generated by deep RNA-sequencing (RNA-seq) of hrSC cultures (n=5) and of neuroblastic tumor cells (NB cells) (n=5), which serve as models for neuronal cells. Comparison of the transcriptomes of hrSCs and NB cells revealed 5822 differentially expressed genes (q-value<0.01, |log2FC>1|), of which 3057 were upregulated and 2754 were down-regulated in hrSCs **(Suppl. Table 2)**. Subsequent functional annotation analysis of genes unique to hrSCs demonstrated gene ontology (GO) terms prominent in MHC class I and class II protein complexes, cellular response to cytokine stimulus, and cytokine mediated signaling pathways **(Fig. 1D, Suppl. Table 3)**. These results were further supported by a gene set enrichment analysis (GSEA), which confirmed the enrichment of genes associated with antigen processing and presentation alongside with cytokine signaling and an inflammatory response in hrSCs **(Fig. 1E, Suppl. Table 4)**. Taken together, transcriptome profiling of hrSCs provides further evidence of cytokine signaling and upregulation of MHCII in response to nerve damage.

### Human repair-related SCs express MHCII and the co-signaling molecules CD40, CD80, CD86, CD58, HVEM and B7-H3

As the primary function of APCs is to modulate T-cell activation through MHCII and the expression of co-signalling molecules, we further investigated whether the latter can be found on hrSCs. Therefore, we cultured hrSCs from eight different donor nerves and used flow cytometry to profile the expression of MHCII and selected co-signalling molecules. As the interaction of SCs with immune cells has been described as a dynamic process (Gold, Zielasek, Kiefer, Toyka, & Hartung, 1996), the analysis was performed at two time points, in passage one and passage two hrSC cultures. SC identity was determined by S100 expression, a well-established SC marker, and showed that mean purity of hrSCs cultures was 82% in passage one (p1) and 70% in passage 2 (p2) **(Fig. 2A)**. The S100 negative cells, presumably nerve-associated fibroblasts, were excluded from further analysis **(Fig. 2A)**. In the S100 positive hrSCs, we quantified the surface expression of MHCII and co-signalling molecules CD40, CD80, CD86, B7-H3, HVEM, PD-L1, PD-L2 and CD58. About 76% and 87% of hrSCs were positive for MHCII in p1 and p2, respectively **(Fig. 2B)**, which is in line with our previously published observation that MHCII expression increased with prolonged culture time (Weiss et al., 2016). Further, p1 as well as p2 hrSCs demonstrated expression of CD40, CD80, B7-H3, CD58, and HVEM **(Fig. 2C-G)**. In contrast, neither PD-L1 nor PD-L2 were detected in p1 or p2 hrSC **(Fig. 2H-I)**. Interestingly, most of p1 hrSCs were negative for CD86, while it was significantly upregulated in p2 cells **(Fig. 2J)**. Hence, hSCs indeed express - next to MHCII - several canonical co-signalling molecules that are required for T-cell activation and inhibition.

**Figure 2.**
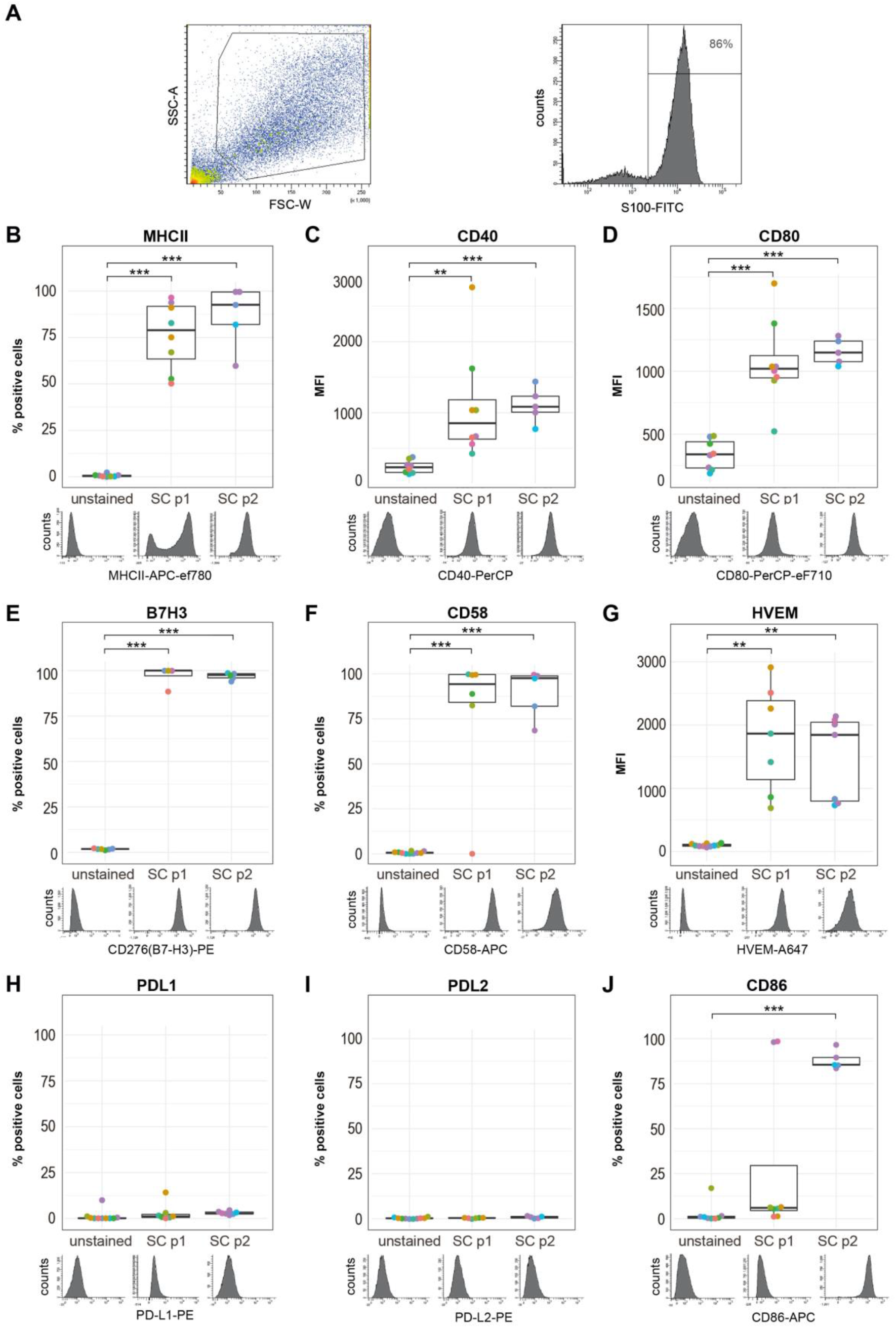
Flow cytometry phenotyping of MHCII and co-signaling molecules present on human repair-related Schwann cells. **(A)** Gating strategy for the identification of S100 positive hrSCs illustrated for one representative experiment. Intact cells are gated in the FSC vs SSC blot and S100 positive cells were selected for further analysis. **(B-J)** Box plots show the expression status of MHCII and co-signaling molecules CD40, CD80, B7H3, CD58, HVEM, PD-L1, PD-L2 and CD86 of S100 positive hrSCs in passage 1 (p1) and p2; technical replicates (same color); biological replicates (different color). The histograms underneath depict one representative experiment. Each biological replicate is conducted with hrSCs isolated from a different donor nerve. **(B, E, F, H, J)** Box plots represent the percentage of positive cells based on gates set in relation to unstained controls as displayed in the histograms. **(C, D, G, I)** Boxplots represent the mean fluorescence intensity (MFI). Boxes contain 50% of data and whiskers the upper and lower 25%; means are displayed as black horizontal lines. A two-way ANOVA using a post-hoc Holm p value correction was performed. * p ≤ 0.05; ** p ≤ 0.01; *** p ≤ 0.001

### The stimulation of TLR3 and TLR4 had no effect on the expression of co-signaling molecules in human repair-related Schwann cells

Professional as well as non-professional APCs can upregulate the expression of co-signaling molecules upon toll-like receptor (TLR) ligation (Chen & Flies, 2013; Fitzgerald & Kagan, 2020; Mehrfeld, Zenner, Kornek, & Lukacs-Kornek, 2018). Our RNA-seq data of hrSCs showed enrichment in TLR signaling in comparison to neuronal cells **(Suppl. Table 4)**, which motivated us to explore the expression of co-signaling molecules in response to inflammatory stimulation. First, we investigated which TLRs are expressed by hrSCs, to choose the corresponding ligands for further analysis. We found elevated expression levels of TLR1, TLR3, TLR4 and TLR6 mRNA **(Fig. 3A, Suppl. Fig. 1)**. As TLR1 and TLR6 mainly function as heterodimers with TLR2, which was not expressed, we focused on TLR3 and TLR4. We thus stimulated p1 hrSC with the TLR4 agonist LPS and TLR3 agonist POLY:IC for 24 hours and subsequently analysed the expression status of co-signalling molecules **(Fig. 3A, Suppl. Fig. 1)**. Interestingly, neither the addition of LPS nor POLY:IC caused a significant expression change of the analyzed co-signaling molecules in hrSCs **(Fig. 3B-E)**, which might be due to the low expression levels of TLR3 and TLR4 **(Fig.3A)**. Upon stimulation with POLY:IC a trend towards upregulation of CD40 and HVEM was seen, but substantial donor variance was observed **(Fig. 3B-E)**. CD40 is not only a co-stimulatory molecule, but also a molecule that facilitates the activation of APCs upon binding of CD40-ligand (Chen & Flies, 2013). CD40-ligand did, however, not affect the expression of CD40 or any other co-signaling molecules tested **(Fig. 3B)**. Together these data show that TLR and CD40 ligation do not affect the expression of co-signaling molecules.

**Figure 3.**
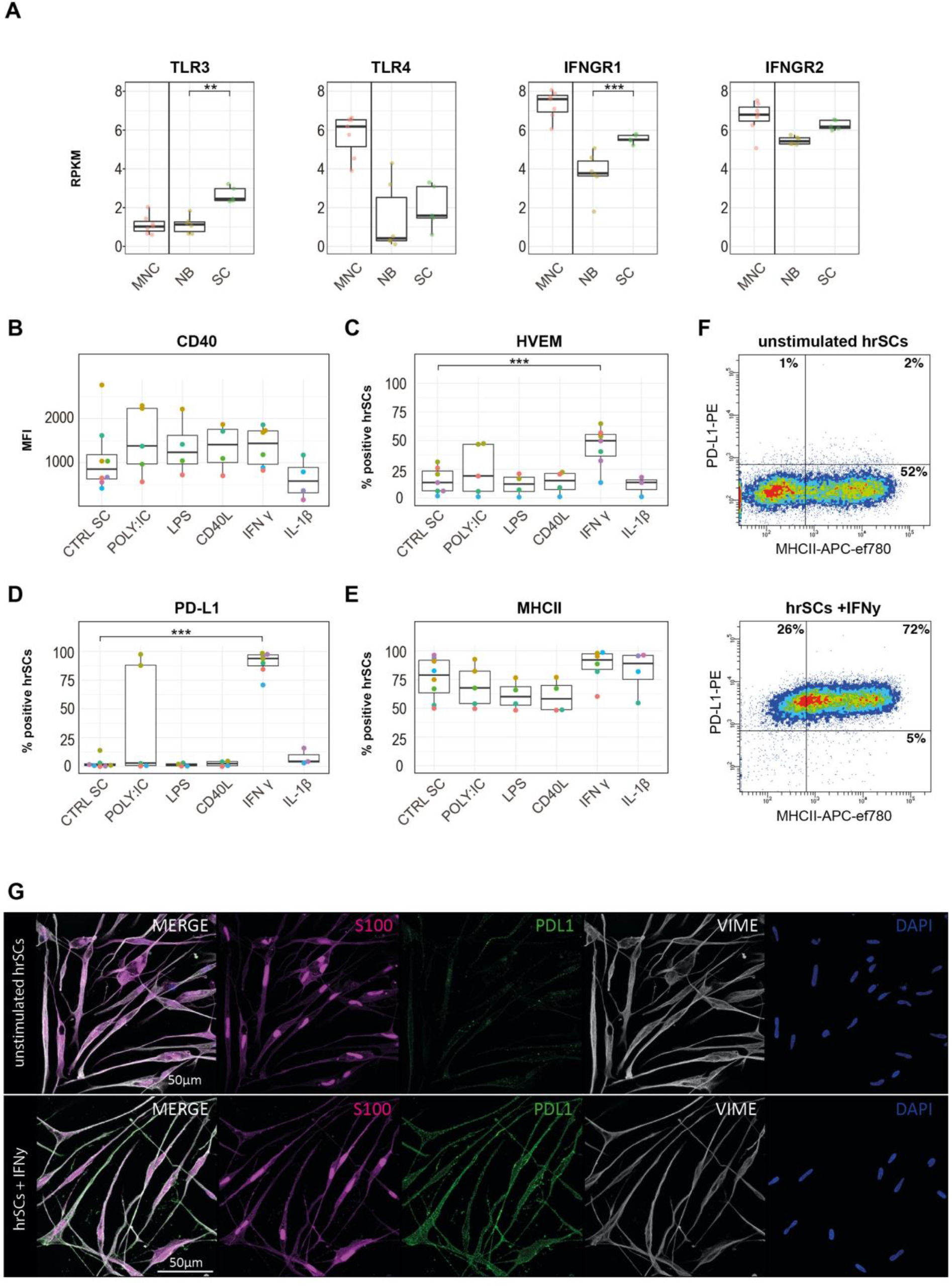
Immunophenotyping of human repair-related Schwann cells upon toll-like receptor and cytokine stimulation. **(A)** Enrichment of toll-like receptor (TLR)-related pathways. GSEA of RNA-seq data sets of hrSCs compared to NB cultures. Box plots show the mRNA expression in reads per kilobase million (RPKM) of TLR3, IFNGR1 and IFNGR2 by RNA-seq in NB cultures (NB, n=5) and hrSCs (SC, n=5). Bone marrow mononuclear cells (MNC, n=5) are shown as reference. **(B-E)** FACS analysis of p1 cultures of hrSCs stimulated with POLY:IC, LPS, CD40L, IFNγ and IL-1β for 24 h. Box plots show the MFI of CD40 (B), percentage of HVEM (C), PD-L1 (D) and MHCII (E) positive hrSCs in p1 cultures stimulated with POLY:IC, LPS, CD40L, IFNγ and IL-1β. Each biological replicate is conducted with hrSCs isolated from a different donor nerve. Boxes contain 50% of data and whiskers the upper and lower 25%. Means are displayed as black horizontal lines. All experiments were performed in at least 4 independent biological replicates. A two-way ANOVA using a post-hoc Holm p value correction was performed; * p ≤ 0.05; ** p ≤ 0.01; *** p ≤ 0.001. **(F)** Representative FACS plots showing PD-L1 vs. MHCII expression of p1 hrSCs either unstimulated (upper plot) and after IFNy stimulation. **(G)** Immunofluorescence image of p1 hrSCs at day 2 after purification without (upper panels) or with (lower panels) IFNy stimulation. HrSC cultures are stained for S100 (magenta), PD-L1 (green), vimentin (grey) and DAPI (blue).

### Human repair-related Schwann cells upregulate HVEM and PD-L1 upon stimulation with IFNγ

Inflammatory processes in peripheral nerve tissues as well as upon injury responses involve the release of pro-inflammatory mediators such as IFNγ and IL-1β by macrophages (Chiu, Von Hehn, & Woolf, 2012; Yao, Graham, Akahata, Oh, & Jacobson, 2010). Notably, we found that the expression of both IFNγ genes, *IFNGR1* and *IFNGR2*, was also upregulated in hrSC in comparison to neuronal cells **(Fig. 3A)**. We further investigated the response of hrSCs to IFNy as well as IL-1β. While IL-1β did not alter the expression of MHCII and any of the co-stimulatory molecules tested, IFNy led to a significant increase in HVEM expression **(Fig. 3C)**. In addition, IFNγ stimulation induced a profound upregulation of PD-L1 **(Fig. 3D)**. PD-L1 expression in response to IFNy was further validated on stimulated hrSC using multicolor immunofluorescence stainings for S100, PD-L1 and vimentin **(Fig. 3G)**. Concordant with our flow cytometry data, unstimulated p1 hrSC did not show a notable PD-L1 staining, whereas IFNγ stimulation strongly induced PD-L1 protein expression **(Fig. 3G)**. These findings show that hrSCs can respond to IFNy, but not to IL-1β, by a significant up-regulation of the co-stimulatory molecule HVEM and the immune check-point molecule PD-L1.

### Secretome analysis of human repair-related Schwann cells reveals a broad spectrum of immunoactive mediators

As the inducible expression of co-signalling molecules by hrSCs together with their well-described function to recruit macrophages and neutrophils (Stratton et al., 2016; Tzekova, Heinen, & Küry, 2014) points towards their active involvement in shaping a local immune response upon nerve injury, we further investigated a panel of immunoregulatory molecules secreted by hrSC. In order to model the *in vivo* situation following nerve injury, supernatants of hrSCs cultured in the absence or presence of neuronal cells and neuronal cell control cultures were analysed. Secretome analysis was performed using a protein array able to detect 274 different secreted factors. A total of 84 secreted molecules were unique to hrSCs and one was only secreted by NB cells (q <0.05). None of the 84 proteins was differentially secreted in the SC-NB co-culture model as compared to SCs alone. We therefore considered these secreted proteins to be derived from hrSCs and further compared those to secreted proteins obtained from NB cultures **(Suppl. Table 5)**. HrSC secreted factors included interleukins IL-6, IL-11, IL-15, TNFα, IFNγ, molecules involved in phagocyte attraction and activation such as MCP-3, MCP-4 or CXCL-16, molecules for neutrophil attraction and activation such as GRO, MIP3-alpha, IL-8 (CXCL-8) or ENA-78 **(Fig. 4A)**. Many of these molecules also directly acting on lymphocytes **(Fig. 4A)**, for example IL-6 that is stimulating the proliferation of antibody producing B-lymphocytes or IL-15 stimualting T-and NK-cell proliferation. Interestingly, hrSCs secreted osteopontin, a molecule involved in multiple processes including the induction of IFNγ through NF-κB activation (Icer & Gezmen-Karadag, 2018). These findings demonstrate the plethora of immunoactive mediators secreted by hrSCs and suggests autocrine activity as well as paracrine modulation of myeloid cells and lymphocytes.

**Figure 4.**
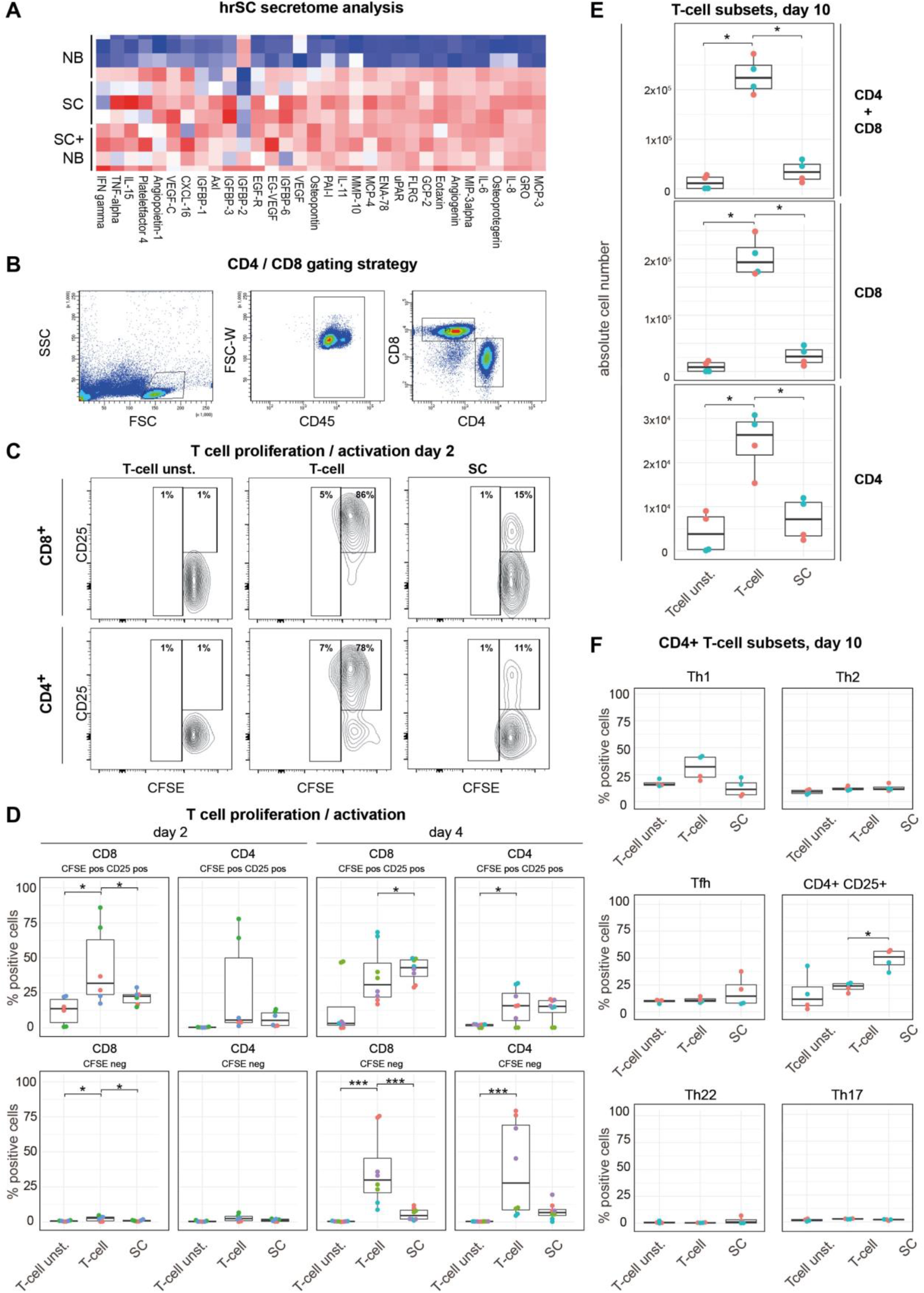
HrSCs secrete immunoactive mediators and inhibit allogenic T-cell activation. **(A)** Secretome analysis by antibody array. Heatmap displays the top 32 differentially secreted proteins (q < 0.05, |log2FC>0.3|) of hrSCs (n=5) and SC-NB co-cultures (n=5) vs. NB cultures (as neuronal cell model) (n=5). (B-F) Allogeneic CD3^+^ T-cells were cultured for 2, 4 or 10 days in the absence (Tcell) or presence of hrSCs (SC) and stimulated with anti-CD3/CD28 beads and analysed by flow cytometry. As control unstimulated T-cells (Tcell unst.) were cultured. **(B)** Representative FACS plots show the gating strategy for CD4^+^ and CD8^+^ T-cells. **(C)** Representative FACS plots showing CD25 expression against CFSE of T-cells at day 2 of co-cultivation and in control cultures. **(D)** Boxplots show the absolute number of CFSE^+^/CD25^+^ and CFSE^−^ CD4^+^ and CD8^+^ T-cells at day 2 and day 4 based on gates set as illustrated in **(B-C)**. **(E)** Boxplots show the absolute number of alive CD4^+^, CD8^+^ or combined CD4^+^and CD8^+^ cells and **(F)** percentage of CD4^+^ subsets evaluated via flow cytometry at day 10 based on gates set as published by (Mahnke et al., 2013). **(D-F)** Boxes contain 50% of data and whiskers the upper and lower 25% Means are displayed as black horizontal lines. All experiments were performed in at least 3 independent biological replicates. A two-way ANOVA using a post-hoc Holm p value correction was performed; * p ≤ 0.05; ** p ≤ 0.01; *** p ≤ 0.001.

### Human repair-related Schwann cells reduce proliferation of allogeneic T-cells

Based on the overlapping features of hrSCs and APCs, i.e. phagocytosis, expression of MHCII, co-signaling and immune checkpoint molecules, and their inducible upregulation and secretion of T-cell modulatory molecules, we next asked whether hrSCs might affect T-cell activation and fate in the nerve injury context. Thus, we performed co-culture assays of hrSCs and allogenic T-cells. More specifically, we used T-cells from healthy donors stimulated with anti-CD3/CD28 beads to simulate an inflammatory environment similarly to peripheral nerve injury. We then evaluated the impact of co-culture assessing the total number of activated (CFSE^+^CD25^+^) and proliferated (CFSE^−/dim^) CD4^+^ and CD8^+^ T-cells **(Fig. 4B-C)**. At day 2 stimulated T-cells showed a significant increase in CFSE^+^CD25^+^ and to a minor extend proliferated (CFSE^−/dim^) CD8^+^ T-cells whereas the presence of hrSCs in co-culture significantly reduced this effect **(Fig. 4D)**. At day 4, stimulation of T-cells resulted in an increase of both, CD8^+^ and CD4^+^, CFSE^+^CD25^+^ as well as proliferated (CFSE^−/dim^) fraction. While proliferation was significantly reduced in CD8^+^ T-cells by the presence of hrSCs in co-culture, CFSE^+^CD25^+^ cells slightly increased **(Fig. 4D)**. A similar trend was observed in the CD4^+^ population **(Fig. 4D)**. This effect became even more apparent at day 10, when the total number of CD8^+^ as well as CD4^+^ T-cells was reduced to numbers comparable with those of unstimulated T-cells. **(Fig. 4E)**. Further, we interrogated the CD4^+^ T-cell population for a potential shift among T-helper subpopulations, i.e. Th1, Th2, Tfh, Th17 and Th22, using a 13-plex flow cytometry panels as previously published by Mahnke et al (Mahnke et al., 2013). Notably, there was no clear trend towards a specific CD4^+^ Th subset, yet a significantly higher percentage of CD4^+^CD25^+^ cells was detected **(Fig. 4F)**. In summary, hrSCs delayed proliferation of CD8^+^ and CD4^+^ T-cells, while promoting the long-term survival or potential switch towards a CD4^+^CD25^+^ phenotype.

## Discussion

Building on our previous characterization of the transcriptome and proteome of human repair-type SCs (Weiss et al., 2016) and studies on SCs and nerve inflammation in rodents (Hartlehnert et al., 2017; Meyer zu Hörste et al., 2010), we aimed to provide novel insight into the immunocompetence of human SCs in an injury condition. Using primary human repair-related SCs, hrSCs, as a unique model to study their immunophenotype and associated functional aspects, our study presents several layers of evidence that hrSCs possess features and functions of APCs and are capable of mediating T-cell dependent immunity. We demonstrate that hrSCs can express CD40, CD80, B7H3, CD58, CD86, HVEM and PD-L1 in addition to MHCII, secrete numerous immunomodulatory molecules, and inhibit allogeneic T-cell activation. It is well accepted that many functions of professional APCs, including the presentation of antigen via MHCII, the expression of co-signalling molecules and the secretion of anti- and pro-inflammatory molecules, are also carried out by non-professional APCs such as mast cells, eosinophils, and non-hematopoietic cells like epithelial cells (Kambayashi & Laufer, 2014; Schuijs, Hammad, & Lambrecht, 2019). With this study, we add compelling evidence that hrSCs could also act as non-professional APCs that may modulate the inflammatory processes within injured nerves. Studying the interaction of primary hrSCs and T-cells allowed the development of a broadly applicable functional *in vitro* model that contributes to the ongoing research in the field of neuroinflammatory disorders, regenerative medicine and immune oncology.

### Human repair-related SCs possess features of antigen presenting cells

In line with our previous studies we show that hrSCs actively perform phagocytosis and express MHCII on their surface (Weiss et al., 2016). It has been recently described that rodent SCs have the ability to regulate MHCII dependent immunity, as the deletion of MHCII lead to a decreased infiltration of CD4+ cells to the site of nerve injury (Hartlehnert et al., 2017). This suggests a functional necessity of MHCII expression in nerve repair. The ability of SCs to present ingested molecules via MHCII and shape an immune response has been suggested before (Baetas-da-Cruz et al., 2009; Meyer Zu Horste et al., 2010; Steinhoff & Kaufmann, 1988; Van Rhijn et al., 2000), yet more detailed studies were needed to understand the role of SCs in phagocytosis, antigen presentation and immune cell modulation in human SCs.

It is well-established that not only classical APCs but also non-professional APCs, like mast cells or epithelial cells, are capable of phagocytosis and antigen presentation via MHC II to CD4+ T-cells (Kambayashi & Laufer, 2014; Schuijs et al., 2019). As non-professional APCs of non-hematopoietic origin do not primarily migrate to lymph nodes, their role in priming naïve T-cells may be less relevant, but their modulation of a local T-cell responses is widely accepted (Kambayashi & Laufer, 2014). Thus, the expression of co-signalling molecules alongside with MHCII and the secretion of other immunomodulatory molecules defines the effect of non-professional APCs in different tissues and conditions. In this study, we show that hrSC are able to express the co-signaling molecules CD58, CD80, and CD86 in addition to MHCII. These molecules are associated with an activation of T-cells (Chen & Flies, 2013; Greenwald, Freeman, & Sharpe, 2004).

Interestingly, a study comparing nerve biopsies of healthy patients and patients with chronic inflammatory demyelinating polyneuropathy (CIDP) identified that CD58 expressing SCs were only found in the latter (Van Rhijn et al., 2000). This could be due to a similarity in the role of SCs after nerve injury and during the interaction with immune cells in autoimmune diseases. However, Van Rhijn et al. did not observe CD86 or CD80 expressing SCs in healthy or CIDP patients, which indicates that the expression of co-signaling molecules detected in our *in vitro* model reflects a unique feature of hrSCs. Of note, our primary human SCs were isolated from peripheral nerves obtained after amputational surgeries and knowledge on pre-existing conditions such as non-diagnosed neuropathies and medication are limited. However, we observed consistent and reproducible effects throughout our molecular, phenotypic and functional characterization of hrSCs. Future studies should combine the knowledge derived from human nerve biopsies and isolated human SCs to validate the immunomodulatory capacities of this unique cell type in nerve injury and different diseases.

In addition to surface expression of co-signalling molecules, we found that hrSC secrete a variety of immunomodulators. This is in line with previous studies on human SCs that demonstrated the secretion of IL-6, IL-8, IL-15 and MCP-1 (Ozaki, Nagai, Lee, Myong, & Kim, 2008; Rutkowski et al., 1999). Our study enriches the repertoire of secreted hrSC molecules by cytokines like IL-11 and chemoattractants like MCP-3, MCP-4, CXCL-16, GRO and MIP3α suggesting an unexpected functional diversity. In contrast to Rutkowski et al., 1999, we did not detect the expression of IL-1β in our assay (Rutkowski et al., 1999). Taken together, these findings demonstrate that hrSCs express the co-signalling molecules CD58, CD80 and CD86 together with MHCII and provide novel insight into the repertoire of hrSC secreted molecules, beyond neurotrophins, with immunoregulatory functions.

### Inhibition of allogeneic T-cell activation - evidence for an immuno-regulatory function of human repair SCs

Furthermore, we show that the exposure to hrSCs causes a delayed or even abrogated CD4^+^ T-cell proliferation and activation. In line with this finding, we demonstrate that SCs express co-inhibitory molecules such as B7-H3, HVEM and provide the first report that hrSC upregulate PD-L1 after stimulation with IFNγ. As not only the activation, but also the termination and resolution of inflammation through surface expression of inhibitory molecules is a hallmark of APCs, the presence of these molecules in hrSCs is remarkable. The source for IFNγ release in inflammatory tissues are mainly NK, CD4^+^ and CD8^+^ T-cells as well as macrophages. HrSCs may even trigger the release of IFNy via secreted osteopontin that has been shown to induce IFNγ production in T-cells (Icer & Gezmen-Karadag, 2018). In this study we show that hrSCs are also capable of IFNγ secretion. Whether the autocrine production of IFNγ by hrSCs or the paracrine IFNγ released by other cell types present at the site of nerve injury induces the surface expression of PD-L1 on hrSCs remains to be determined.

We further observed that delayed T-cell activation was not accompanied by a shift towards any particular T helper subset, but rather resulted in CD4+ T-cells with high CD25 expression, which may represent a regulatory or exhausted phenotype. The hypothesis of regulatory/exhausted T-cells is supported by previous observations in rodent models (Meyer zu Horste et al., 2014; Meyer zu Hörste et al., 2010; F.-J. Wang, Cui, & Qian, 2018; X. Wang et al., 2014). Similarly, it has been shown that non-professional APCs like type II alveolar epithelial cells can prime antigen specific CD4+ T-cells towards regulatory T-cells (Kambayashi & Laufer, 2014). This suggests that hrSCs in their activated state might, despite high MHCII, CD40, CD80 and CD58 expression and secretion of pro-inflammatory cytokines IL-6, IL-8, TNFα and IFNγ, adopt an antigen presenting cell phenotype similar to M2 macrophages, which tightly control and terminate T-cell responses via B7-H3 and the PD-L1/PD-1 axis. Thus, a balance between the initiation and termination of an immune response may be essentially controlled through repair SCs during the multistep process of nerve regeneration.

The expression of PD-L1 by hrSCs after stimulation with IFNγ supports the idea of a time and situation dependent role of repair SCs. It is tempting to speculate that repair SCs initiate the termination of the inflammatory response they helped to induce as first responders to nerve injury. Our data suggest that repair SCs might possess an immunoregulatory function that could prevent unnecessary damage to the neuronal environment by terminating an exceeding immune response, in which large amounts of IFNγ are produced by immune cells recruited to the site of injury. To address this possibility, deeper phenotypic and functional characterization of especially CD4+CD25+ T-cells *in vitro/ex vivo* will be required in the future. Furthermore, manipulating the balance of pro- and anti-inflammatory profile of repair SCs might represent a novel therapeutic target in regenerative and pathological processes.

### Implications for the field of neuro-inflammatory disorders, regenerative medicine and immune oncology

As the interaction between SCs and immune cells is of importance not only after nerve injury but also regarding infectious, inflammatory and autoimmune disease of the peripheral nervous system, a comprehensive understanding of this interaction is crucial. It has recently been suggested that the activation of T-helper cells via MHCII by SC promotes neuropathic pain and axonal loss after nerve injury in mice (Hartlehnert et al., 2017). In this regard the presented panel of co-signaling molecules provides additional pharmaceutical targets to tackle this interaction. In addition, it has been shown that rats with experimental autoimmune neuritis, a common model for Guillain-Barrè syndrome, showed a clinical improvement, reduced neuronal lymphocyte infiltration and a shift towards regulatory T-cells in the peripheral blood after administration of PD-L1 (Ding et al., 2016).

Importantly, it could be shown that SCs play a fundamental role in certain immune-oncological processes. SCs with a repair-related phenotype (including MHCII expression) are attracted by favorable forms of peripheral neuroblastic tumors, neuroblastomas, of a genetic subtype and trigger tumor cell maturation/differentiation and apoptosis, a phenomenon which could also be recapitulated in *in vitro* experiments (Weiss et al., 2021). These tumors, in comparison to their malignant counterpart, frequently show prominent MHCII+ and CD3+ immune cell infiltrates (Ambros et al., 1996; Weiss et al., 2021). It will be interesting to study the composition of these infiltrates and whether and how SCs contribute to their recruitment and modulation.

In summary, we here provide *in vitro* evidence that human SCs in an injury condition adopt functions of APCs, i.e, phagocytosis, up-regulation of MHCII and co-signaling molecules, secretion of an array of immunoregulatory molecules, and repression of T-cell activation. Our data suggest that repair SCs can participate in the termination of the inflammatory response to prevent excessive tissue damage and allow nerve regeneration. The molecules expressed and secreted by hrSC presented in this study will help to understand their complex interplay with immune cells after injury and in disease.

## Author contributions

S.T.-M. conceptualized the project; J.B., T.W. and S.T.-M. planned experiments, performed research, analyzed and interpreted data and wrote the manuscript; H.S. and F.R. performed research and analyzed data; M.K. developed bioinformatics tools and analyzed data; A.D. and P.S. provided essential reagents, planned experiments and interpreted data; R.W. provided essential material; P.F.A. and I.M.A interpreted data; all authors reviewed the manuscript.

## Acknowledgements

This study was supported by the Herzfeldersche Familienstiftung (Grants to S. Taschner-Mandl), Modicell (MC-IAPP Project 285875, to S. Taschner-Mandl), FFG Visiomics (Project 10959423 to S. Taschner-Mandl) and FWF Liquidhope (Project I4162 to S. Taschner-Mandl) and St. Anna Kinderkrebsforschung. We are also grateful to Dieter Printz and the FACS core facility (Children’s Cancer Research Institute) for excellent technical support.

## Conflict of Interest Statement

The authors declare no conflict of interest.

## Supplementary Figures

**Supplementary Figure 1.**
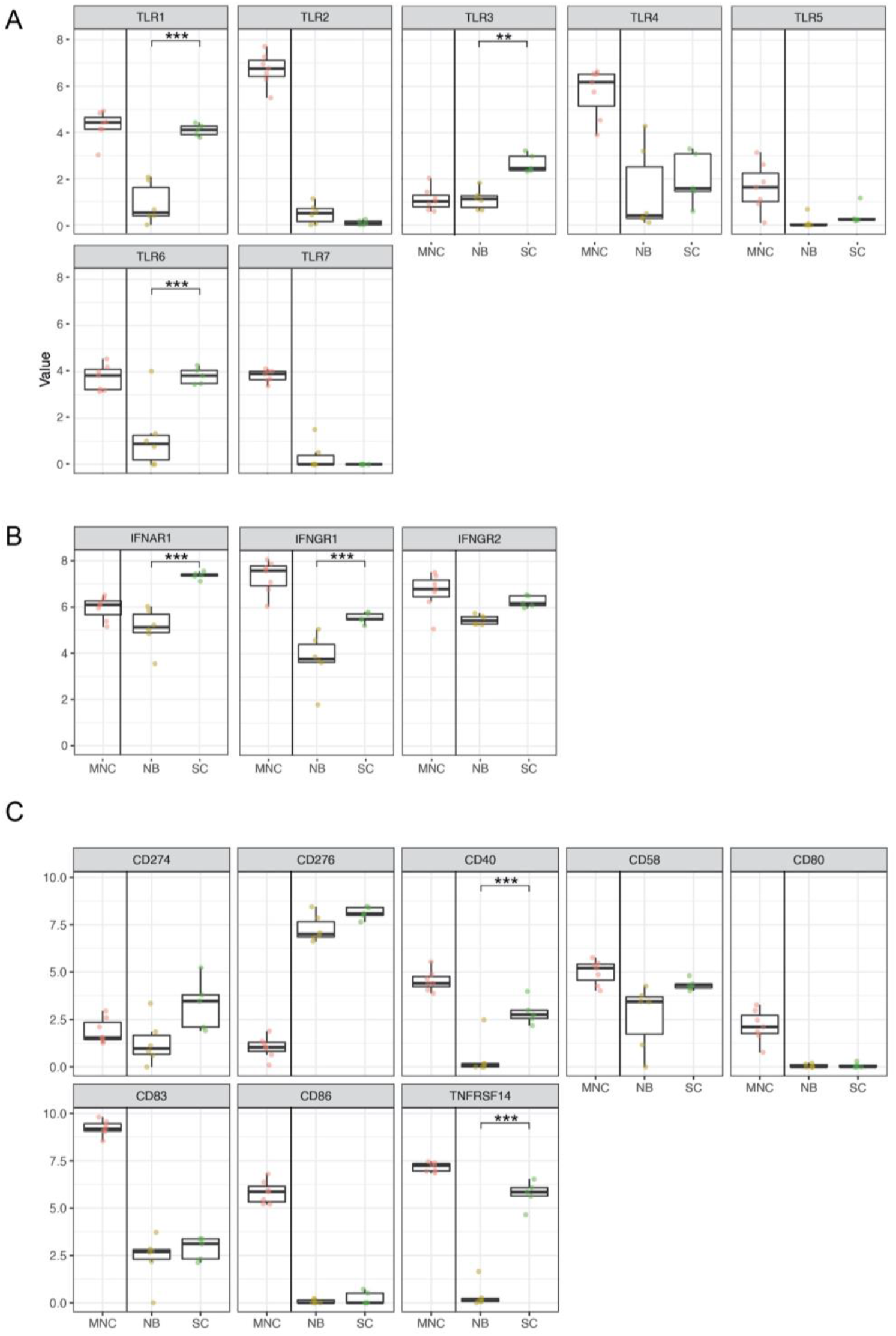
mRNA expression of Toll-like receptors, interferon receptors and co-signaling molecules. Boxplots show mRNA levels (RPKM) in hrSCs (n=5) versus NB cells (n=5). MNCs are shown as reference. **(A)** Toll-like receptors **(B)** interferon receptors and **(C)** co-stimulatory and –inhibitory molecules. Boxes contain 50% of data and whiskers the upper and lower 25% means are displayed as black horizontal lines.

**Supplementary Figure 2.**
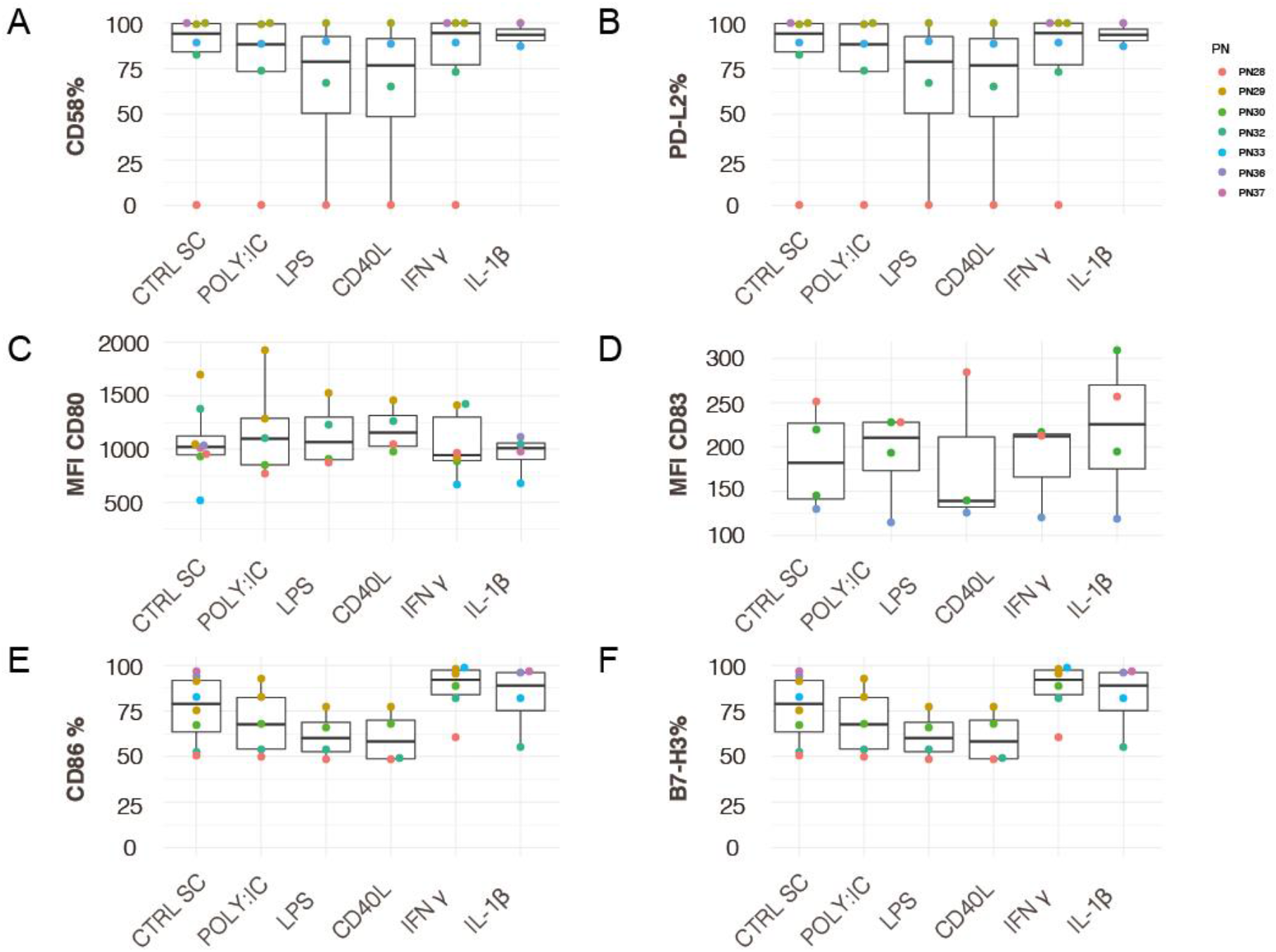
Flow cytometry-based phenotyping of MHCII and co-signaling molecules upon inflammatory stimulation. Box plots show the expression status of CD58 **(A)**, PDL2 **(B)**, CD80 **(C)**, CD83 **(D)**, CD86 **(E)** and B7H3 **(F)** of S100 positive hrSCs after stimulation with POLY:IC, LPS, CD40L, IFNγ and IL-1β; n=9; technical replicates (same color); biological replicates (different color). Each biological replicate is conducted with hrSCs isolated from a different donor nerve. **(A, B, E, F)** Boxplots represent the percentage of positive cells based on gates set in relation to unstained controls. **(C, D)** Boxplots represent the mean fluorescence intensity (MFI). Boxes contain 50% of data and whiskers the upper and lower 25%; means are displayed as black horizontal lines. A two-way ANOVA using a post-hoc Holm p value correction was performed; * p ≤ 0.05; ** p ≤ 0.01; *** p ≤ 0.001.

## Supplementary Tables

**Supplementary Table 1.** List of antibodies. perm = permeabilization necessary, RT = room temperature.

| 1<sup>st</sup> antibodies |  |  |  | Application |  |
| --- | --- | --- | --- | --- | --- |
| Antigen | Species | catalog No | company | dilution | comment |
| S100 | rabbit | #Z0311 | DAKO | 1:200 | 1 hr, RT, perm |
| vimentin | chicken | #AB5733 | Merck Millipore | 1:200 | 1 hr, RT, perm |
| NGFR | rabbit | #8238S | CellSignaling | 1:300 | o.n., 4°C |
| PD-L1 | Mouse | #4274539 | eBioscience | 1:50 | o.n. 4°C |
| HLA-DR-α1 | mouse | #M0746 | DAKO | 1:50 | o.n., 4°C |

| 2<sup>nd</sup> antibodies |  |  |  | Application |  |
| --- | --- | --- | --- | --- | --- |
| Antigen | Species | catalog No | company | dilution | comment |
| α rb FITC | swine | #F0205 | DAKO | 1:50 | 1 hr, RT |
| α ch AF647 | goat | #SA5-10073 | LifeTech. | 1:300 | 1 hr, RT |
| α ms AF594 | goat | #A11032 | LifeTech. | 1:300 | 1 hr, RT |

Supplementary Table 1 | List of antibodies.
| directly labelled antibodies |  |  |  | Application |  |
| --- | --- | --- | --- | --- | --- |
| Antigen | Species | catalog No | company | dilution | comment |
| S100B-FITC* | rabbit | #Z0311 | DAKO | 1:50 | 20 min, 4°C, perm |
| CD80-PerCP-eF710 | mouse | 46-0809-42 | eBioscience | 1:25 |  |
| CD276(B7-H3)-PE | mouse | 565829 | BD Bioscience | 1:50 |  |
| CD40-PerCP | mouse | Ab91282 | Abcam | 1:5 |  |
| MHCII-APC-ef780 | mouse | 47-9956-42 | eBioscience | 1:50 |  |
| PD-L1-PE | mouse | 557924 | BD Bioscience | 1:5 |  |
| HVEM-A647 | mouse | 564411 | BD Bioscience | 1:25 |  |
| CD86-APC | mouse | 555660 | BD Bioscience | 1:25 |  |
| CD273- PE | mouse | 565829 | BD Bioscience | 1:25 |  |
| CD58-APC | mouse | 17-0578-41 | eBioscience | 1:25 |  |

**Supplementary Table 2 | RNA-sequencing. Differential gene expression analysis of hrSCs compared to NB cell lines.**

**Supplementary Table 3 | RNA-sequencing. GO functional annotation of top 250 genes upregulated in hrSCs versus NB cell lines.**

**Supplementary Table 4 | RNA-sequencing. Gene set enrichment analysis (GSEA) in hrSC versus NB cell lines.**

**Supplementary Table 5 | Protein array. Differential protein secretion of hrSCs versus NB cell lines.**

## Notes

### Competing Interest Statement

The authors have declared no competing interest.

## References

Ambros, I M, Rumpler, S., Luegmayr, A., Hattinger, C. M., Strehl, S., Kovar, H., & Ambros, P. F. (1997). Neuroblastoma cells can actively eliminate supernumerary MYCN gene copies by micronucleus formation--sign of tumour cell revertance? European Journal of Cancer (Oxford, England : 1990), 33(12), 2043–2049. https://doi.org/10.1016/s0959-8049(97)00204-9

Ambros, Ingeborg M., Zellner, A., Roald, B., Amann, G., Ladenstein, R., Printz, D., & Ambros, P. F. (1996). Role of Ploidy, Chromosome 1p, and Schwann Cells in the Maturation of Neuroblastoma. New England Journal of Medicine, 334(23), 1505–1511. https://doi.org/10.1056/NEJM199606063342304

Armati, P. J., Pollard, J. D., & Gatenby, P. (1990). Rat and human Schwann cells in vitro can synthesize and express MHC molecules. Muscle & Nerve, 13(2), 106–116. https://doi.org/10.1002/mus.880130204

Azam, S. H., & Pecot, C. V. (2016). Cancer’s got nerve: Schwann cells drive perineural invasion. The Journal of Clinical Investigation, 126(4), 1242–1244. https://doi.org/10.1172/JCI86801

Baetas-da-Cruz, W., Alves, L., Pessolani, M. C. V, Barbosa, H. S., Régnier-Vigouroux, A., Corte-Real, S., & Cavalcante, L. a. (2009). Schwann cells express the macrophage mannose receptor and MHC class II. Do they have a role in antigen presentation? Journal of the Peripheral Nervous System : JPNS, 14(2), 84–92. https://doi.org/10.1111/j.1529-8027.2009.00217.x

Bergsteinsdottir, K., Kingston, A., Mirsky, R., & Jessen, K. R. (1991). Rat Schwann cells produce interleukin-1. Journal of Neuroimmunology, 34(1), 15–23. https://doi.org/10.1016/0165-5728(91)90094-N

Biedler, J. L., Helson, L., & Spengler, B. A. (1973). Morphology and growth, tumorigenicity, and cytogenetics of human neuroblastoma cells in continuous culture. Cancer Research, 33(11), 2643–2652.

Biedler, J. L., Roffler-Tarlov, S., Schachner, M., & Freedman, L. S. (1978). Multiple neurotransmitter synthesis by human neuroblastoma cell lines and clones. Cancer Research, 38(11 Pt 1), 3751–3757.

Bunimovich, Y. L., Keskinov, A. A., Shurin, G. V, & Shurin, M. R. (2017). Schwann cells: a new player in the tumor microenvironment. Cancer Immunology, Immunotherapy : CII, 66(8), 959–968. https://doi.org/10.1007/s00262-016-1929-z

Chen, L., & Flies, D. B. (2013). Molecular mechanisms of T cell co-stimulation and co-inhibition. Nature Reviews. Immunology, 13(4), 227–242. https://doi.org/10.1038/nri3405

Chiu, I. M., Von Hehn, C. A., & Woolf, C. J. (2012). Neurogenic inflammation and the peripheral nervous system in host defense and immunopathology. Nature Neuroscience. https://doi.org/10.1038/nn.3144

Combaret, V., Turc-Carel, C., Thiesse, P., Rebillard, A. C., Frappaz, D., Haus, O., & Favrot, M. C. (1995). Sensitive detection of numerical and structural aberrations of chromosome 1 in neuroblastoma by interphase fluorescence in situ hybridization. Comparison with restriction fragment length polymorphism and conventional cytogenetic analyses. International Journal of Cancer, 61(2), 185–191. https://doi.org/10.1002/ijc.2910610208

Crawford, S. E., Stellmach, V., Ranalli, M., Huang, X., Huang, L., Volpert, O., & Bouck, N. (2001). Pigment epithelium-derived factor (PEDF) in neuroblastoma: a multifunctional mediator of Schwann cell antitumor activity. Journal of Cell Science, 114(Pt 24), 4421–4428.

Ding, Y., Han, R., Jiang, W., Xiao, J., Liu, H., Chen, X., & Hao, J. (2016). Programmed Death Ligand 1 Plays a Neuroprotective Role in Experimental Autoimmune Neuritis by Controlling Peripheral Nervous System Inflammation of Rats. Journal of Immunology (Baltimore, Md. : 1950), 197(10), 3831–3840. https://doi.org/10.4049/jimmunol.1601083

Direder, M., Weiss, T., Copic, D., Vorstandlechner, V., Laggner, M., Mildner, C. S., & Mildner, M. (2021). Schwann cells contribute to keloid formation. MedRxiv, 2021.08.09.21261701. https://doi.org/10.1101/2021.08.09.21261701

Dobin, A., Davis, C. A., Schlesinger, F., Drenkow, J., Zaleski, C., Jha, S., & Gingeras, T. R. (2013). STAR: ultrafast universal RNA-seq aligner. Bioinformatics (Oxford, England), 29(1), 15–21. https://doi.org/10.1093/bioinformatics/bts635

Duan, R.-S., Jin, T., Yang, X., Mix, E., Adem, A., & Zhu, J. (2007). Apolipoprotein E deficiency enhances the antigen-presenting capacity of Schwann cells. Glia, 55(7), 772–776. https://doi.org/10.1002/glia.20498

Fischer, M., & Berthold, F. (2003). Characterization of the gene expression profile of neuroblastoma cell line IMR-5 using serial analysis of gene expression. Cancer Letters, 190(1), 79–87. https://doi.org/10.1016/s0304-3835(02)00581-5

Fitzgerald, K. A., & Kagan, J. C. (2020). Toll-like Receptors and the Control of Immunity. Cell, 180(6), 1044–1066. https://doi.org/10.1016/j.cell.2020.02.041

Gaudet, A. D., Popovich, P. G., & Ramer, M. S. (2011). Wallerian degeneration: gaining perspective on inflammatory events after peripheral nerve injury. Journal of Neuroinflammation, 8, 110. https://doi.org/10.1186/1742-2094-8-110

Gentleman, R. C., Carey, V. J., Bates, D. M., Bolstad, B., Dettling, M., Dudoit, S., & Zhang, J. (2004). Bioconductor: open software development for computational biology and bioinformatics. Genome Biology, 5(10), R80. https://doi.org/10.1186/gb-2004-5-10-r80

Goethals, S., Ydens, E., Timmerman, V., & Janssens, S. (2010). Toll-like receptor expression in the peripheral nerve. GLIA, 58(14), 1701–1709. https://doi.org/10.1002/glia.21041

Gold, R., Zielasek, J., Kiefer, R., Toyka, K. V, & Hartung, H. P. (1996). Secretion of nitrite by Schwann cells and its effect on T-cell activation in vitro. Cellular Immunology, 168(1), 69–77. https://doi.org/10.1006/cimm.1996.0050

Gomez-Sanchez, J. A., Carty, L., Iruarrizaga-Lejarreta, M., Palomo-Irigoyen, M., Varela-Rey, M., Griffith, M., & Jessen, K. R. (2015). Schwann cell autophagy, myelinophagy, initiates myelin clearance from injured nerves. The Journal of Cell Biology, 210(1), 153–168. https://doi.org/10.1083/jcb.201503019

Greenwald, R. J., Freeman, G. J., & Sharpe, A. H. (2004). THE B7 FAMILY REVISITED. Annual Review of Immunology, 23(1), 515–548. https://doi.org/10.1146/annurev.immunol.23.021704.115611

Hartlehnert, M., Derksen, A., Hagenacker, T., Kindermann, D., Schäfers, M., Pawlak, M., & Meyer zu Horste, G. (2017). Schwann cells promote post-traumatic nerve inflammation and neuropathic pain through MHC class II. Scientific Reports. London. https://doi.org/10.1038/s41598-017-12744-2

Hartley, S. W., & Mullikin, J. C. (2015). QoRTs: a comprehensive toolset for quality control and data processing of RNA-Seq experiments. BMC Bioinformatics, 16(1), 224. https://doi.org/10.1186/s12859-015-0670-5

Hörste, G. M. Z., Hu, W., Hartung, H. P., Lehmann, H. C., & Kieseier, B. C. (2008). The immunocompetence of Schwann cells. Muscle and Nerve. https://doi.org/10.1002/mus.20893

Icer, M. A., & Gezmen-Karadag, M. (2018). The multiple functions and mechanisms of osteopontin. Clinical Biochemistry, 59, 17–24. https://doi.org/10.1016/j.clinbiochem.2018.07.003

Jang, S. Y., Shin, Y. K., Park, S. Y., Park, J. Y., Lee, H. J., Yoo, Y. H., & Park, H. T. (2016). Autophagic myelin destruction by Schwann cells during Wallerian degeneration and segmental demyelination. Glia, 64(5), 730–742. https://doi.org/10.1002/glia.22957

Jessen, K. R., & Mirsky, R. (2016). The repair Schwann cell and its function in regenerating nerves. The Journal of Physiology, 594(13), 3521–3531. https://doi.org/10.1113/JP270874

Kaisho, T., & Akira, S. (2000). Critical roles of Toll-like receptors in host defense. Critical Reviews in Immunology, 20(5), 393–405.

Kambayashi, T., & Laufer, T. M. (2014). Atypical MHC class II-expressing antigen-presenting cells: can anything replace a dendritic cell? Nature Reviews. Immunology, 14(11), 719–730. https://doi.org/10.1038/nri3754

Karanth, S., Yang, G., Yeh, J., & Richardson, P. M. (2006). Nature of signals that initiate the immune response during Wallerian degeneration of peripheral nerves. Experimental Neurology, 202(1), 161–166. https://doi.org/10.1016/j.expneurol.2006.05.024

Kingston, A. E., Bergsteinsdottir, K., Jessen, K. R., Van der Meide, P. H., Colston, M. J., & Mirsky, R. (1989). Schwann cells co-cultured with stimulated T cells and antigen express major histocompatibility complex (MHC) class II determinants without interferon-gamma pretreatment: synergistic effects of interferon-gamma and tumor necrosis factor on MHC class II in. European Journal of Immunology, 19(1), 177–183. https://doi.org/10.1002/eji.1830190128

La Fleur, M., Underwood, J. L., Rappolee, D. A., & Werb, Z. (1996). Basement membrane and repair of injury to peripheral nerve: defining a potential role for macrophages, matrix metalloproteinases, and tissue inhibitor of metalloproteinases-1. The Journal of Experimental Medicine, 184(6), 2311–2326. https://doi.org/10.1084/jem.184.6.2311

Law, C. W., Chen, Y., Shi, W., & Smyth, G. K. (2014). voom: Precision weights unlock linear model analysis tools for RNA-seq read counts. Genome Biology, 15(2), R29. https://doi.org/10.1186/gb-2014-15-2-r29

Lee, H., Jo, E. K., Choi, S. Y., Oh, S. B., Park, K., Soo Kim, J., & Lee, S. J. (2006). Necrotic neuronal cells induce inflammatory Schwann cell activation via TLR2 and TLR3: Implication in Wallerian degeneration. Biochemical and Biophysical Research Communications, 350(3), 742–747. https://doi.org/10.1016/j.bbrc.2006.09.108

Lilje, O., & Armati, P. J. (1997). The distribution and abundance of MHC and ICAM-1 on Schwann cells in vitro. Journal of Neuroimmunology, 77(1), 75–84. https://doi.org/10.1016/S0165-5728(97)00063-5

Mahnke, Y. D., Beddall, M. H., & Roederer, M. (2013). OMIP-017: Human CD4+ helper T-cell subsets including follicular helper cells. Cytometry Part A, 83 A(5), 439–440. https://doi.org/10.1002/cyto.a.22269

Mancardi, G. L., Cadoni, A., Zicca, A., Schenone, A., Tabaton, M., De Martini, I., & Zaccheo, D. (1988). HLA-DR Schwann cell reactivity in peripheral neuropathies of different origins. Neurology, 38(6), 848–851. https://doi.org/10.1212/wnl.38.6.848

Mehrfeld, C., Zenner, S., Kornek, M., & Lukacs-Kornek, V. (2018). The Contribution of Non-Professional Antigen-Presenting Cells to Immunity and Tolerance in the Liver. Frontiers in Immunology, 9, 635. https://doi.org/10.3389/fimmu.2018.00635

Meyer zu Horste, G., Cordes, S., Mausberg, A. K., Zozulya, A. L., Wessig, C., Sparwasser, T., & Kieseier, B. C. (2014). FoxP3+ regulatory T cells determine disease severity in rodent models of inflammatory neuropathies. PloS One, 9(10), e108756. https://doi.org/10.1371/journal.pone.0108756

Meyer Zu Horste, G., Heidenreich, H., Lehmann, H. C., Ferrone, S., Hartung, H.-P., Wiendl, H., & Kieseier, B. C. (2010). Expression of antigen processing and presenting molecules by Schwann cells in inflammatory neuropathies. Glia, 58(1), 80–92. https://doi.org/10.1002/glia.20903

Meyer zu Hörste, G., Heidenreich, H., Mausberg, A. K., Lehmann, H. C., ten Asbroek, A. L. M. A., Saavedra, J. T., & Kieseier, B. C. (2010). Mouse Schwann cells activate MHC class I and II restricted T-cell responses, but require external peptide processing for MHC class II presentation. Neurobiology of Disease, 37(2), 483–490. https://doi.org/10.1016/j.nbd.2009.11.006

Meyer zu Hörste, G., Hu, W., Hartung, H.-P., Lehmann, H. C., & Kieseier, B. C. (2008). The immunocompetence of Schwann cells. Muscle & Nerve, 37(1), 3–13. https://doi.org/10.1002/mus.20893

Momoi, M., Kennett, R. H., & Glick, M. C. (1980). A membrane glycoprotein from human neuroblastoma cells isolated with the use of a monoclonal antibody. The Journal of Biological Chemistry, 255(24), 11914–11921.

Mootha, V. K., Lindgren, C. M., Eriksson, K.-F., Subramanian, A., Sihag, S., Lehar, J., & Groop, L. C. (2003). PGC-1α-responsive genes involved in oxidative phosphorylation are coordinately downregulated in human diabetes. Nature Genetics, 34(3), 267–273. https://doi.org/10.1038/ng1180

Murata, K.-Y., & Dalakas, M. C. (2000). Expression of the co-stimulatory molecule BB-1, the ligands CTLA-4 and CD28 and their mRNAs in chronic inflammatory demyelinating polyneuropathy. Brain : A Journal of Neurology, 123 (Pt 8, 1660–1666. https://doi.org/10.1016/S0002-9440(10)65141-3

Nocera, G., & Jacob, C. (2020). Mechanisms of Schwann cell plasticity involved in peripheral nerve repair after injury. Cellular and Molecular Life Sciences : CMLS, 77(20), 3977–3989. https://doi.org/10.1007/s00018-020-03516-9

Ozaki, A., Nagai, A., Lee, Y. B., Myong, N. H., & Kim, S. U. (2008). Expression of cytokines and cytokine receptors in human Schwann cells. Neuroreport, 19(1), 31–35. https://doi.org/10.1097/WNR.0b013e3282f27e60

Ritchie, M. E., Phipson, B., Wu, D., Hu, Y., Law, C. W., Shi, W., & Smyth, G. K. (2015). limma powers differential expression analyses for RNA-sequencing and microarray studies. Nucleic Acids Research, 43(7), e47. https://doi.org/10.1093/nar/gkv007

Roberts, S. L., Dun, X.-P., Doddrell, R. D. S., Mindos, T., Drake, L. K., Onaitis, M. W., & Parkinson, D. B. (2017). Sox2 expression in Schwann cells inhibits myelination in vivo and induces influx of macrophages to the nerve. Development (Cambridge, England), 144(17), 3114–3125. https://doi.org/10.1242/dev.150656

Rutkowski, J. L., Tuite, G. F., Lincoln, P. M., Boyer, P. J., Tennekoon, G. I., & Kunkel, S. L. (1999). Signals for proinflammatory cytokine secretion by human Schwann cells. Journal of Neuroimmunology, 101(1), 47–60. https://doi.org/10.1016/s0165-5728(99)00132-0

Samuel, N. M., Mirsky, R., Grange, J. M., & Jessen, K. R. (1987). Expression of major histocompatibility complex class I and class II antigens in human Schwann cell cultures and effects of infection with Mycobacterium leprae. Clinical and Experimental Immunology, 68(3), 500–509.

Schuijs, M. J., Hammad, H., & Lambrecht, B. N. (2019). Professional and “Amateur” Antigen-Presenting Cells In Type 2 Immunity. Trends in Immunology, 40(1), 22–34. https://doi.org/10.1016/j.it.2018.11.001

Spierings, E., De Boer, T., Zulianello, L., & Ottenhoff, T. H. M. (2000). Novel mechanisms in the immunopathogenesis of leprosy nerve damage: The role of schwann cells, T cells and Mycobacterium leprae. Immunology and Cell Biology. https://doi.org/10.1046/j.1440-1711.2000.00939.x

Steinhoff, U., & Kaufmann, S. H. (1988). Specific lysis by CD8+ T cells of Schwann cells expressing Mycobacterium leprae antigens. European Journal of Immunology, 18(6), 969–972. https://doi.org/10.1002/eji.1830180622

Stock, C., Bozsaky, E., Watzinger, F., Poetschger, U., Orel, L., Lion, T., & Ambros, P. F. (2008). Genes proximal and distal to MYCN are highly expressed in human neuroblastoma as visualized by comparative expressed sequence hybridization. The American Journal of Pathology, 172(1), 203–214. https://doi.org/10.2353/ajpath.2008.061263

Stratton, J. A., & Shah, P. T. (2016). Macrophage polarization in nerve injury: do Schwann cells play a role? Neural Regeneration Research, 11(1), 53–57. https://doi.org/10.4103/1673-5374.175042

Stratton, J. A., Shah, P. T., Kumar, R., Stykel, M. G., Shapira, Y., Grochmal, J., & Midha, R. (2016). The immunomodulatory properties of adult skin-derived precursor Schwann cells: implications for peripheral nerve injury therapy. The European Journal of Neuroscience, 43(3), 365–375. https://doi.org/10.1111/ejn.13006

Subramanian, A., Tamayo, P., Mootha, V. K., Mukherjee, S., Ebert, B. L., Gillette, M. A., & Mesirov, J. P. (2005). Gene set enrichment analysis: A knowledge-based approach for interpreting genome-wide expression profiles. Proceedings of the National Academy of Sciences, 102(43), 15545 LP–15550. https://doi.org/10.1073/pnas.0506580102

Toews, A. D., Barrett, C., & Morell, P. (1998). Monocyte chemoattractant protein 1 is responsible for macrophage recruitment following injury to sciatic nerve. Journal of Neuroscience Research, 53(2), 260–267. https://doi.org/10.1002/(SICI)1097-4547(19980715)53:2<260::AID-JNR15>3.0.CO;2-A

Tofaris, G. K., Patterson, P. H., Jessen, K. R., & Mirsky, R. (2002). Denervated Schwann cells attract macrophages by secretion of leukemia inhibitory factor (LIF) and monocyte chemoattractant protein-1 in a process regulated by interleukin-6 and LIF. The Journal of Neuroscience : The Official Journal of the Society for Neuroscience, 22(15), 6696–6703. https://doi.org/10.1523/JNEUROSCI.22-15-06696.2002

Tzekova, N., Heinen, A., & Küry, P. (2014). Molecules involved in the crosstalk between immune- and peripheral nerve Schwann cells. Journal of Clinical Immunology. https://doi.org/10.1007/s10875-014-0015-6

Van Rhijn, I., Van den Berg, L. H., Bosboom, W. M. J., Otten, H. G., & Logtenberg, T. (2000). Expression of accessory molecules for T-cell activation in peripheral nerve of patients with CIDP and vasculitic neuropathy. Brain, 123(10), 2020–2029. https://doi.org/10.1093/brain/123.10.2020

Wang, F.-J., Cui, D., & Qian, W.-D. (2018). Therapeutic Effect of CD4+CD25+ Regulatory T Cells Amplified In Vitro on Experimental Autoimmune Neuritis in Rats. Cellular Physiology and Biochemistry : International Journal of Experimental Cellular Physiology, Biochemistry, and Pharmacology, 47(1), 390–402. https://doi.org/10.1159/000489919

Wang, X., Zheng, X.-Y., Ma, C., Wang, X.-K., Wu, J., Adem, A., & Zhang, H.-L. (2014). Mitigated Tregs and augmented Th17 cells and cytokines are associated with severity of experimental autoimmune neuritis. Scandinavian Journal of Immunology, 80(3), 180–190. https://doi.org/10.1111/sji.12201

Weiss, T., Taschner-Mandl, S., Ambros, P. F., & Ambros, I. M. (2018). Detailed Protocols for the Isolation, Culture, Enrichment and Immunostaining of Primary Human Schwann Cells. Methods in Molecular Biology (Clifton, N.J.), 1739, 67–86. https://doi.org/10.1007/978-1-4939-7649-2_5

Weiss, T., Taschner-Mandl, S., Bileck, A., Slany, A., Kromp, F., Rifatbegovic, F., & Ambros, I. M. (2016). Proteomics and transcriptomics of peripheral nerve tissue and cells unravel new aspects of the human Schwann cell repair phenotype. GLIA, 64(12), 2133–2153. https://doi.org/10.1002/glia.23045

Weiss, T., Taschner-Mandl, S., Janker, L., Bileck, A., Rifatbegovic, F., Kromp, F., & Ambros, I. M. (2021). Schwann cell plasticity regulates neuroblastic tumor cell differentiation via epidermal growth factor-like protein 8. Nature Communications, 12(1), 1624. https://doi.org/10.1038/s41467-021-21859-0

Wekerle, H., Schwab, M., Linington, C., & Meyermann, R. (1986). Antigen presentation in the peripheral nervous system: Schwann cells present endogenous myelin autoantigens to lymphocytes. European Journal of Immunology, 16(12), 1551–1557. https://doi.org/10.1002/eji.1830161214

Yao, K., Graham, J., Akahata, Y., Oh, U., & Jacobson, S. (2010). Mechanism of neuroinflammation: enhanced cytotoxicity and IL-17 production via CD46 binding. Journal of Neuroimmune Pharmacology : The Official Journal of the Society on NeuroImmune Pharmacology, 5(3), 469–478. https://doi.org/10.1007/s11481-010-9232-9

Ydens, E., Cauwels, A., Asselbergh, B., Goethals, S., Peeraer, L., Lornet, G., & Janssens, S. (2012). Acute injury in the peripheral nervous system triggers an alternative macrophage response. Journal of Neuroinflammation, 9, 176. https://doi.org/10.1186/1742-2094-9-176

Ydens, E., Lornet, G., Smits, V., Goethals, S., Timmerman, V., & Janssens, S. (2013). The neuroinflammatory role of Schwann cells in disease. Neurobiology of Disease, 55, 95–103. https://doi.org/10.1016/j.nbd.2013.03.005

Zhang, S. H., Shurin, G. V, Khosravi, H., Kazi, R., Kruglov, O., Shurin, M. R., & Bunimovich, Y. L. (2020). Immunomodulation by Schwann cells in disease. Cancer Immunology, Immunotherapy : CII, 69(2), 245–253. https://doi.org/10.1007/s00262-019-02424-7

